# Resonance modulation, annihilation and generation of antiresonance and antiphasonance in 3D neuronal systems: interplay of resonant and amplifying currents with slow dynamics

**DOI:** 10.1101/091207

**Authors:** Horacio G. Rotstein

**Affiliations:** Department of Mathematical Sciences New Jersey Institute of Technology Newark, NJ 07102, USA

## Abstract

Subthreshold (membrane potential) resonance and phasonance (preferred amplitude and zero-phase responses to oscillatory inputs) in single neurons arise from the interaction between positive and negative feedback effects provided by relatively fast amplifying currents and slower resonant currents. In 2D neuronal systems, amplifying currents are required to be slaved to voltage (instantaneously fast) for these phenomena to occur. In higher dimensional systems, additional currents operating at various effective time scales may modulate and annihilate existing resonances and generate antiresonance (minimum amplitude response) and antiphasonance (zero-phase response with phase monotonic properties opposite to phasonance). We use mathematical modeling, numerical simulations and dynamical systems tools to investigate the mechanisms underlying these phenomena in 3D linear models, which are obtained as the linearization of biophysical (conductance-based) models. We characterize the parameter regimes for which the system exhibits the various types of behavior mentioned above in the rather general case in which the underlying 2D system exhibits resonance. We consider two cases: (i) the interplay of two resonant gating variables, and (ii) the interplay of one resonant and one amplifying gating variables. Increasing levels of an amplifying current cause (i) a response amplification if the amplifying current is faster than the resonant current, (ii) resonance and phasonance attenuation and annihilation if the amplifying and resonant currents have identical dynamics, and (iii) antiresonance and antiphasonance if the amplifying current is slower than the resonant current. We investigate the underlying mechanisms by extending the envelope-plane diagram approach developed in previous work (for 2D systems) to three dimensions to include the additional gating variable, and constructing the corresponding envelope curves in these envelope-space diagrams. We find that antiresonance and antiphasonance emerge as the result of an asymptotic boundary layer problem in the frequency domain created by the different balances between the intrinsic time constants of the cell and the input frequency *f* as it changes. For large enough values of *f* the envelope curves are quasi-2D and the impedance profile decrease with the input frequency. In contrast, for *f* ≪ 1 the dynamics is quasi-1D and the impedance profile increases above the limiting value in the other regime. Antiresonance is created because the continuity of the solution requires the impedance profile to connect the portions belonging to the two regimes. If in doing so the phase profile crosses the zero value, then antiphasonance is also generated.

## 1 Introduction

The response of a neuron to oscillatory inputs can be characterized by the so-called impedance (*Z*) and phase (*ϕ*) profiles, which are the curves of the impedance amplitude (or simply impedance) and phase-shift (or simply phase) as a function of the input frequency (*f*) respectively [1–3]. A neuron exhibits subthreshold (membrane potential) resonance and phasonance if the impedance profile peaks at a nonzero input (resonant) frequency *f*_*res*_ and the phase profile vanishes at a non-zero input (phasonant) frequency *f*_*phas*_. The latter indicates that the input and output peak at the same time for *f* = *f*_*phas*_. Several neuron types have been shown to exhibit subthreshold resonance and phasonance in response to oscillatory inputs in both current and voltage clamp experiments and in models [1, 2, 4–40]. Subthreshold resonance, phasonance and intrinsic oscillations are related, but different phenomena, and neurons may exhibit one(s) in the absence of the other(s) [2–4, 41, 42].

The functionality of neuronal resonance has not been fully established yet. However, several lines of work have demonstrated the role of resonance for network oscillatory activity either directly or indirectly [38, 43–47] both experimentally and theoretically. The role played by resonance in network oscillations is further supported by the fact that the resonant frequency of certain neuron types coincides with the oscillatory frequency of the networks in which they are embedded [12, 13, 26, 48]

Subthreshold resonance results from the interaction between positive and negative feedback effects provided by the amplifying and resonant gating variables that govern the dynamics of the participating ionic currents. Neurons with only a passive leak current are low-pass filters: the impedance profile is a decreasing function of *f* and the response is always delayed. Resonance requires the presence of a gating variable that opposes changes in voltage such as these associated to the so-called resonant currents (e.g., hyperpolarization-activated mixed-cation *I*_*h*_ and slow-potassium *I*_*Ks*_). The gating variables associated to so-called amplifying currents (e.g., persistent sodium *I*_*Nap*_ and inward-rectifying potassium *I*_*Kir*_) favor changes in voltage and therefore enhance the voltage response in the resonant frequency band [1–3].

In previous work we have carried out a thorough analysis of subthreshold resonance and phasonance in 2D linearized and nonlinear neuronal models of quadratic type having a fast (instantaneous) amplifying current and a slower resonant current [3, 41, 42]. By combining biophysical (conductance-based) modeling, numerical simulations and dynamical systems tools we have identified the biophysical and dynamic mechanisms underlying the generation of resonance and phasonance and described the effects that changes in the maximal conductances and other model parameters have on *f*_*res*_, *f*_*phas*_ and the additional attributes characterizing the shapes of the impedance and phase profiles. In these studies we have used linear and quadratic models, representing the linearization and “quadratization”, respectively, of biophysical models of Hodgkin-Huxley type [49] describing the neuronal subthreshold dynamics, and we have investigated the role played by various types of resonant and amplifying currents, and their combinations, through these simplified models.

For 2D models to exhibit resonance the fast amplifying gating variable is constrained to be instantaneously fast and slaved to voltage. These models fail to capture complex phenomena such as the presence of troughs in the impedance profile that have been experimentally observed in hippocampal interneurons [18] and more recently in the so-called LP neurons of the crab stomatogastric ganglion pyloric network [37], and have been shown theoretically to emerge in models with slower amplifying gating variables [2]. We refer to this phenomenon as antiresonance and the associated emergence of an additional zero in the phase profile as antiphasonance, and we refer to the corresponding characteristic frequencies as *f*_*ares*_ and *f*_*aphas*_, respectively. For antiresonance and antiphasonance to occur the additional degree of freedom provided by the slower amplifying gating variable is necessary. To our knowledge, the mechanisms of generation of antiresonance and antiphasonance as well as the mechanisms that govern the modulation of underlying 2D resonances by the interaction between two ionic currents with slow dynamics and either the similar or opposite feedback effects in oscillatory forced systems have not been investigated.

The goal of this paper is to address these issues for 3D linear models representing the linearization of conductance-based models. These models have a resonant gating variable and an additional gating variable that can be either resonant or amplifying. This gives rise to two types of interactions between cooperative and competitive feedback effects operating at comparable or different time scales. Antiresonance and antiphasonance occur when the amplifying gating variable is slower than the resonant one. The antiresonant patterns, in the presence of an underlying 2D resonance, are shaped by a trade-off between the time constant and linearized conductance of this amplifying gating variable. In [50] we have investigated how these different ways of interaction shape the intrinsic subthreshold oscillatory properties of linearized and nonlinear models and the complex dependence of these oscillatory patterns on the model parameters. Here we extend our study to models receiving oscillatory forcing. As for 2D systems, this extension is not straightforward, primarily because even linear systems can exhibit subthreshold resonance in the absence of intrinsic subthreshold oscillations [2, 3, 41, 42].

For our mechanistic analysis we extend the envelope-plane diagram approach developed in [41] (see also [42]) to the 3D space to include the additional gating variable. We view the dynamics of the oscillatory forced 3D system as the result of the response limit cycle trajectories tracking the oscillatory motion of the voltage nullsurface as time progresses. This allows us to construct the so-called envelope curves in the 3D envelope-space diagram. The envelope curves consist of the points corresponding to the peaks of these response limit cycle trajectories in the *v*-direction (maximum voltage and the corresponding values of the other coordinates). The envelope-space diagrams contain geometric and dynamic information about the frequency response properties of the system to oscillatory inputs, and are the frequency analogous to the phase-space diagrams. The envelope curves are trajectories in the envelope-space diagrams parametrized by the input frequency as trajectories in the phase-space diagrams are curves parametrized by time. This approach allow us to identify the principles that govern (i) the modulation and annihilation of existing resonances, and (ii) the generation of antiresonance and antiphasonance. In addition, it allows us to determine the conditions under which these phenomena occur. For visualization purposes, we use projections of the envelope-space diagrams onto the relevant planes for the participating variables in addition to the 3D diagrams.

An important outcome of this analysis is the identification of the mechanism of generation of antiresonance and antiphasonance as an asymptotic boundary layer problem in the frequency domain. This is the result of the different balances between the input frequency *f* and the cell’s intrinsic time constants (independent of *f*) as *f* changes. For values of *f* away from zero the dynamics are 2D and governed by the underlying 2D resonant system independently of the slow amplifying variable. In contrast, for values of *f* close to zero the dynamics are quasi-1D and the resistance *Z*(0), which depends on the maximal conductance of the slow variable, increases above the limit of the impedance for the 2D system as *f* approaches zero. The trough in the impedance profile is created because the continuity of the solution requires the impedance profile to connect the portions belonging to the two regimes. If in doing so the phase profile crosses the zero value, then antiphasonance is also generated.

The investigation of the voltage response to sinusoidal inputs has typically focused on how preferred frequency responses are generated by the presence of a negative feedback (resonance and phasonance) and how the voltage response is enhanced at the preferred frequency band by the presence of a positive feedback (amplification). While these two processes are often discussed separately [1, 2], they are intertwined: amplifying currents cannot generate resonance in the absence of a resonant current, but they can modify the values the resonant and phasonant frequencies as well as other attributes of the impedance and phase profiles [3, 41, 51]. Therefore, the study presented here includes the mechanisms of selection of the resonant and phasonant frequencies in addition to the generation and amplification of resonance.

The outline of the paper is as follows. In Section 3.1 we revisit the biophysical and dynamic mechanisms of generation of resonance and phasonance in 2D models (voltage *v* and a resonant gating variable *x*_1_). For future use, we introduce the description of so-called dynamic phase-plane for 2D oscillatory forced systems, which include the envelope-plane diagrams and envelope-curves mentioned above. In Section 3.2 we discuss the three limiting and special cases where the voltage response dynamics of the 3D system (*v, x*_1_ and a second gating variable *x*_2_) reduces to the voltage response dynamics for quasi-2D systems, and therefore they can be understood in terms of the results from previous work. In Section 3.3 we identify the basic mechanisms of modulation and annihilation of resonance / phasonance and the generation of antiresonance / antiphasonance in the forced 3D model. As a baseline case for our study we use a representative set of parameter values for the underlying 2D system such that this 2D system exhibits resonance but no intrinsic oscillations. In Section 3.4 we extend this investigation to understand how the trade-off between the maximal conductance and time constant of the *x*_2_ shape the preferred response properties of the 3D model identified in Section 3.3 using the same baseline parameter values as we used there. We consider the interplay of both (i) two resonant gating variables, and (ii) one resonant and one amplifying gating variables. In Section 3.5 we investigate the dynamic mechanisms of resonance modulation and annihilation and the generation of antiresonance and antiphasonance. To this end, we extend the dynamic phase-space approach discussed in Section 3.1 to include the three variables (voltage and the two gating variables). The mechanistic principles we extract from this study have a more general validity than the specific sets of parameter values considered in the previous sections. Finally, in Section 4 we discuss our results, limitations and implications for neuronal dynamics.

## 2 Methods

### 2.1 Conductance-based models

We use the following biophysical (conductance-based) models of Hodgkin-Huxley type [49, 52] to describe the neuronal subthreshold dynamics

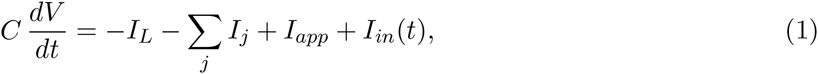

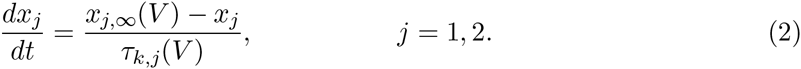

In the current-balance equation (1), *V* is the membrane potential (mV), *t* is time (ms), *C* is the membrane capacitance (*μ*F/cm^2^), *I*_*app*_ is the applied bias (DC) current (*μ*A/cm^2^), *I*_*L*_ = *G*_*L*_ (*V* − *E*_*L*_) is the leak current, and *I*_*j*_ = *G*_*j*_ *x*_*j*_, (*V* − *E*_*j*_) are generic expressions for ionic currents (with *j* an index) with maximal conductance *G*_*j*_ (mS/cm^2^) and reversal potentials *E*_*j*_ (mV) respectively. The dynamics of the gating variables *x*_*j*_ are governed by the kinetic equations (2) where *x*_*j*,∞_(*V*) and *τ*_*j*,*x*_(*V*) are the voltage-dependent activation/inactivation curves and time constants respectively. The generic ionic currents *I*_*j*_ we consider here are restricted to have a single gating variable *x*_*j*_ and to be linear in *x*_*j*_. This is typically the case for persistent sodium (*I*_*Nap*_), *h*- (hyperpolarization-activated, mixed-cation, inward), and slow-potassium (M-type) (*I*_*Ks*_) currents. Our discussion and results can be easily adapted to include ionic currents having two gating variables raised to powers different from one such as T-type calcium and A-type potassium currents [1].

The input current *I*_*in*_(*t*) (*μ*A/cm^2^) in eq. (1) has the form

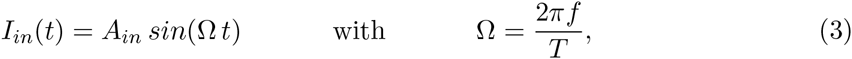

where *T* = 1000 ms and *f* is the input frequency (Hz).

In this paper we focus on 3D models describing the dynamics of *V* and the two gating variables. Additional currents whose gating variables evolve on a very fast time scale (as compared to the other variables) can be included by using the adiabatic approximation *x*_*k*_ = *x*_*k*,∞_(*V*). Here we include one such fast current *I*_3_ = *G*_3_ *x*_3,∞_(*V*) (*V* − *E*_3_). Additional fast currents can be included without significantly changing the formalism used here.

### 2.2 Linearized conductance-based models

Linearization of the autonomous part (*I*_*in*_(*t*) = 0) of system (1)-(2) around the fixed-point (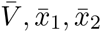) yields [2]

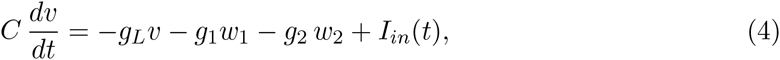

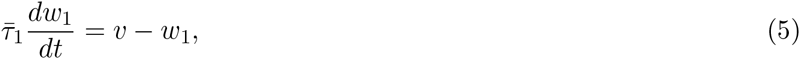

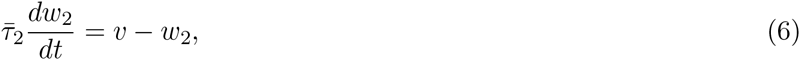

where

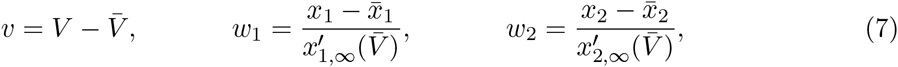

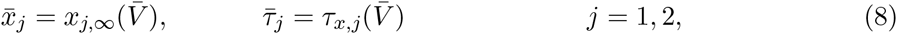

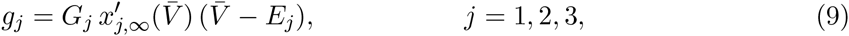

and

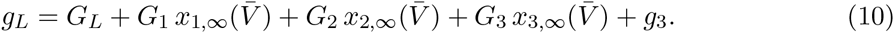

Note that the gating variables *w*_1_ and *w*_2_ in (7) have units of voltage ([*v*] = [*w*_1_] = V).

The effective leak conductance *g*_*L*_ (10) contains information about the biophysical leak conductance *G*_*L*_, the ionic conductances, and their associated voltage-dependent activation / inactivation curves. The fast ionic current *I*_3_ contributes to *g*_*L*_ with an additional term *g*_3_. The signs of the effective ionic conductances *g*_*j*_ determine whether the associated gating variables are either resonant (*g*_*j*_ > 0) or amplifying (*g*_*j*_ < 0) [1, 2]. Specific examples are the gating variables associated to *I*_*h*_ (resonant), *I*_*Ks*_ (resonant), *I*_*Nap*_ (amplifying) and *I*_*Kir*_ (amplifying). All terms in *g*_*L*_ are positive except for the last one that can be either positive or negative. Specifically, *g*_*L*_ can become negative for negative enough values of *g*_3_. This and the other linearized conductances are affected not only by the respective biophysical conductances, but also by the magnitudes and signs of the activation (*σ* < 0) and inactivation (*σ* > 0) curves and their derivatives, which are typically given by expressions of the form

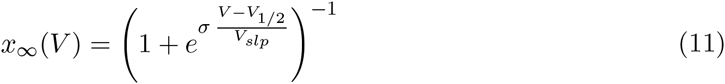

and

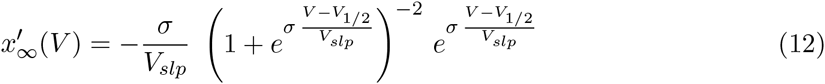

where *V*_1/2_ and *V*_*slp*_ > 0 are constants.

### 2.3 Biophysical- and geometric-based rescaled models

System (4)-(6) can be rescaled in order to reduce the number of parameters of the autonomous part (from six to four) and to capture the relative effects of the linearized conductances *g*_1_ and *g*_2_ and their time constants 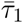 and 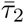. Here we use two different rescalings. The so-called biophysical rescaling uses dimensionless conductances and is appropriate for the analysis of the dependence of the resonant properties in terms of the ionic conductances and the role of the time constants in effectively strengthening these conductances. It is close to the one that combines the biophysical description with the time-scale separation between variables and has been used in [2] (see also [3]). The so-called geometric rescaling is amenable for the mechanistic analysis using dynamical systems tools (“dynamic” phase-space analysis) and is an extension of the one we used in [41]. For the biophysically related questions we will use the original linearized (4)-(6) model. However, the biophysical rescaling shows that the principles extracted from our study are valid beyond the representative parameter values we used.

#### 2.3.1 Rescaled model I: Biophysical dimensionless conductances

Define the following dimensionless time

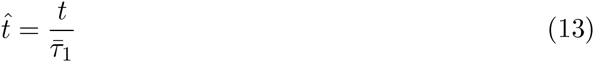

and parameters

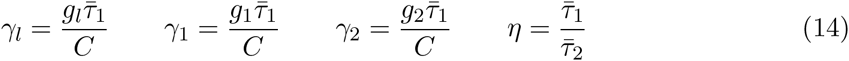

Substitutying into (4)-(5) we obtain

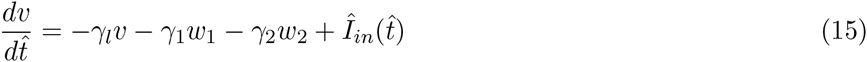

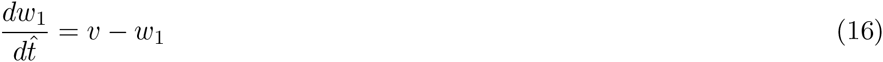

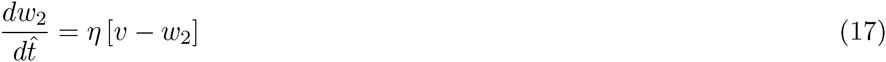

where

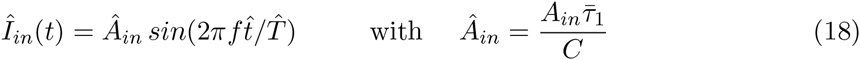

with 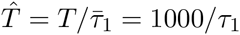. Note that [*f*] = Hz, and [*v*] = [*w*_1_] = [*w*_2_] = V.

#### 2.3.2 Rescaled model II: Geometric/dynamic description

Define the following dimensionless time and (dimensional) voltage variables

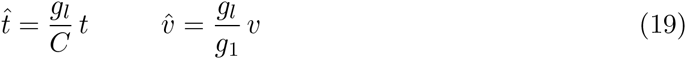

and parameters

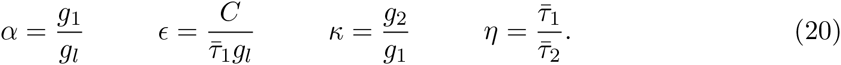

Substitutying into (4)-(5) we obtain

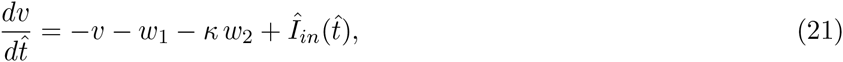

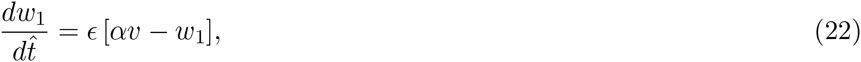

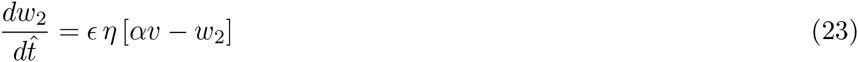

where

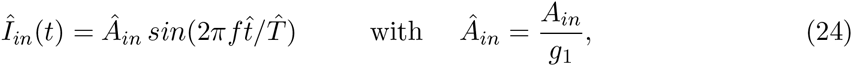

with 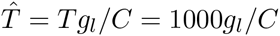. Note that [*f*] = Hz, and [*v*] = [*w*_1_] = [*w*_2_] = V.

System (21)-(23) can be thought of as a 3D extension of the 2D linear system investigated in [41] to which it reduces for *κ* = 0. Geometrically, the parameter *α* is the slope of the *w*-nullcline and can be thought of as representing the strength of the gain of the feedback in the linearized system. The parameter *ϵ* represents the time scale separation between *v* and *w*_1_ and *η* represents the time scale separation between *w*_1_ and *w*_2_.

Since for resonant gating variables *g*_1_ > 0, the sign of both *α* and *ϵ* depends on whether *g*_*L*_ is positive or negative. In the absence of fast amplifying currents (*G*_3_ = *g*_3_ = 0), *g*_*L*_ > 0 and then both *α* > 0 and *ϵ* > 0. When an amplifying current is present and its contribution to *g*_*L*_ is small enough, the sign of both *α* and *ϵ* remains positive. However, when stronger contributions of the fast amplifying current causes *g*_*L*_ to be negative, the sign of both *α* and *ϵ* are also negative. Since resonance becomes amplified as *g*_*L*_ decreases [3], we expect resonance to be more amplified for negative values of both *ϵ* and *α* as compared to positive ones as in [41]. The cases including values of *α* and *ϵ* having different signs are excluded from this study since the underlying autonomous system is either unstable (saddle) or stable (node) but does not exhibit resonance [41].

#### 2.3.3 Linking the rescalings I and II

The parameters in the autonomous parts of the rescaled systems (15)-(17) and (21)-(23) are related by the following equations

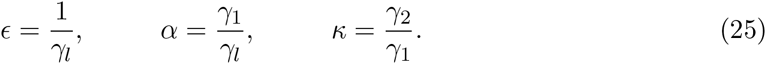

The dimensionless parameter *η* is common to both rescaled models.

### 2.4 Decoupling the input frequency from the oscillatory input

For the type of analysis we present in this paper it is useful to rescale time *t* → *t*/*f* in order to separate the effect of the input frequency *f* from the input’s time dependence [42]. After dropping the “hat” from the time and input current, system (4)-(6) becomes

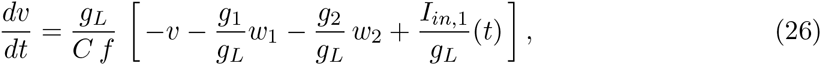

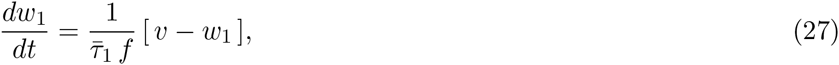

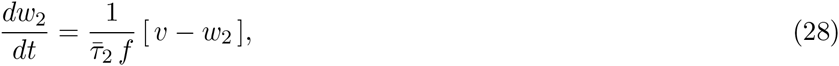

where

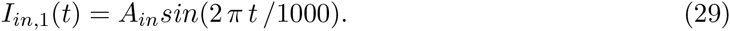

In system (26)-(28) the sinusoidal input function has the same frequency for all values of *f*, which affects the speed of the voltage response. Similarly, system (21)-(23) becomes

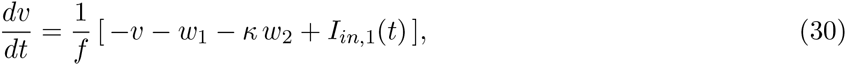

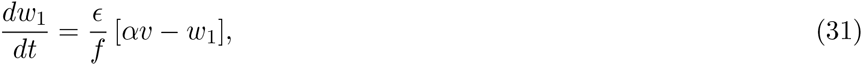

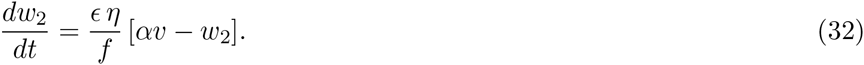

In geometric / dynamic terms, this shows that the input frequency *f* affects the speed of the trajectories in the phase space without affecting the direction of the underlying vector field [41, 42]. Additionally, this transformation highlights the different balances that arise between the effective time constant (*f*^−1^) “imposed” by the oscillatory input and the cell’s intrinsic time constants as the input frequency changes.

When *w*_1_ and *w*_2_ are identical (and have the same initial conditions) the 3D system (30)-(32) is effectively 2D. By substituting these variables by *w* eqs. (30)-(32) reduce to

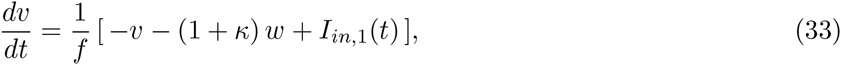

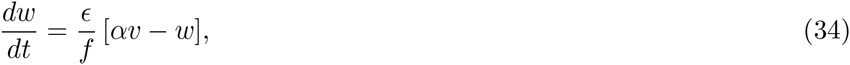

In what follows we drop the “bar” sign from the time constants *τ*_1_ and *τ*_2_.

### 2.5 Voltage response to sinusoidal input currents: impedance amplitude and phase

The voltage response of a neuron receiving an oscillatory current input at a frequency *f* can be characterized by its amplitude (normalized by the input amplitude) and phase (or phase-shift). Together, these quantities constitute the so called impedance function *Z*(*f*) (a complex quantity). For simplicity, here we use the term impedance, and we use the notation *Z*(*f*), to refer to the impedance amplitude. We refer to the graphs of *Z*(*f*) and *ϕ*(*f*) as the impedance and phase profiles.

For a linear system receiving sinusoidal input currents of the form (3), the voltage response is given by

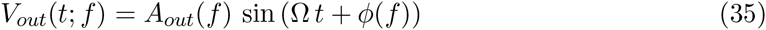

where *A*_*out*_(*f*) is the voltage amplitude. The impedance amplitude is given by

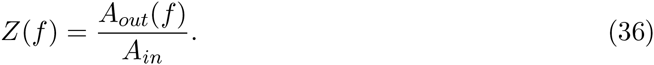

When the input and output frequencies coincide, as it always happens for linear systems, the phase *ϕ*(*f*) captures the difference between the peaks of the voltage output and input current normalized by the oscillation period. Eqs. (57)-(58) in the Appendix give the analytical expressions for the impedance and phase profiles.

#### 2.5.1 Impedance and phase profiles for 2D linear systems: resonance and phasonance

For 2D linear systems the impedance profiles are either a decreasing function of *f* (red curve in Fig. 1-a1), representing a low-pass filter response, or a non-monotonic graph exhibiting a peak *Z*_*max*_ at a non-zero input frequency (blue curve in Fig. 1-a1) representing a resonant response at the (resonant) frequency *f*_*res*_. The phase profile may be either an increasing function of *f* converging asymptotically to *ϕ* = π/2 (red curve in Fig. 1-a2), and representing a delayed response for all values of *f*, or a non-monotonic graph exhibiting a zero-phase response at a non-zero input frequency *f*_*phas*_ (blue curve in Fig. 1-a2) that we refer to as the phasonant frequency. For *f* = *f*_*phas*_ the input current and output voltage peak at the same time. For *f* < *f*_*phas*_, the voltage response is advanced, while for *f* > *f*_*phas*_ the voltage response is delayed. We have investigated the mechanisms of generation of resonance and phasonance in 2D linear systems in [41].

**Figure 1:**
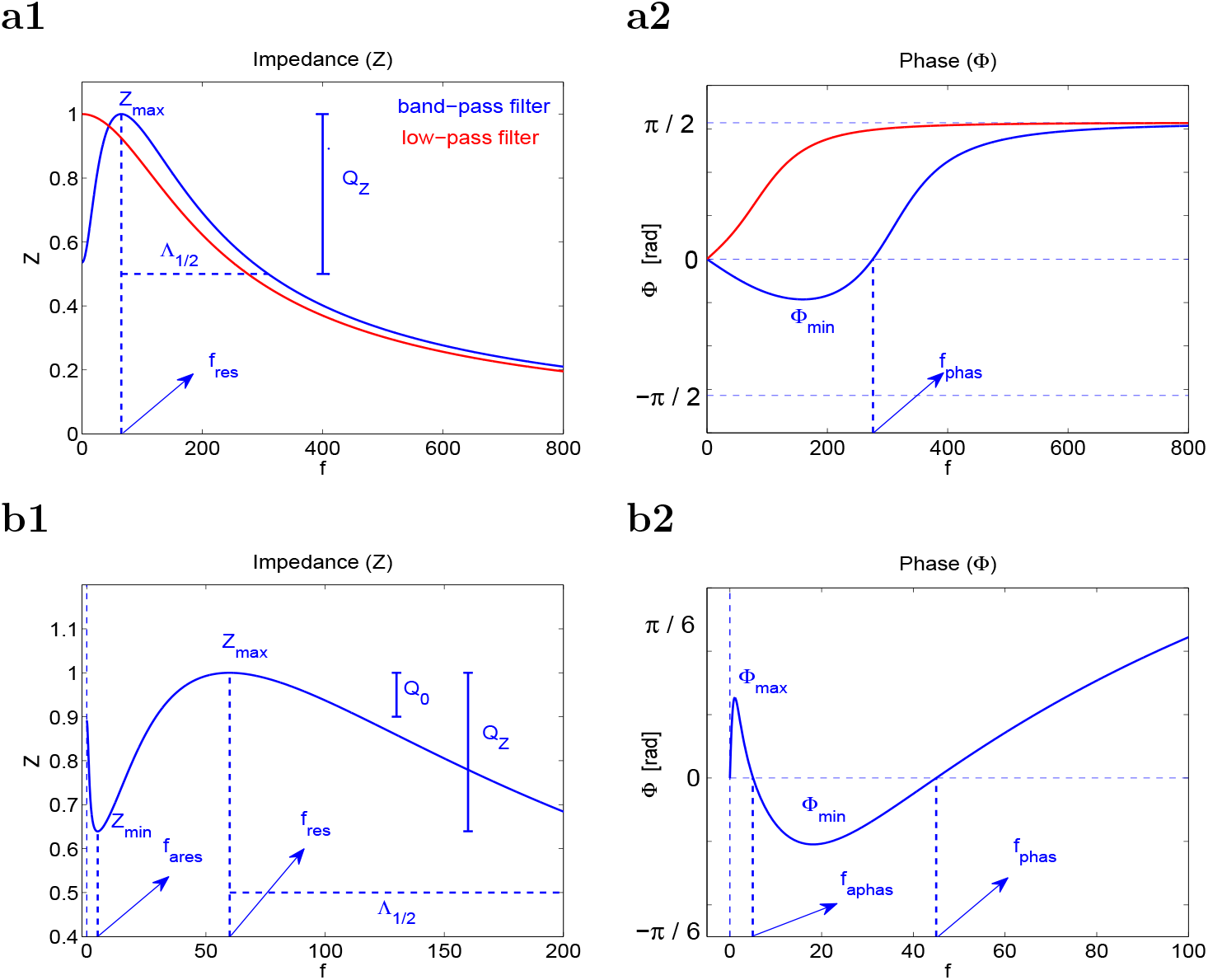
**Representative Impedance (*Z*) and phase (*ϕ*) profiles (curves of *Z* and *ϕ* vs. the input frequency *f*) for 2D (a) and 3D (a,b) linear systems.** (**a1**) The impedance *Z* is characterized by four attributes: the resonant frequency *f*_*res*_, the impedance peak *Z*_*max*_, the resonance amplitude *Q*_*Z*_ = *Z*_*max*_ − *Z*(0), and the half-bandwidth Λ_1/2_. (**a2**) The phase *ϕ* is characterized by two attributes: the zero-crossing frequency *f*_*phas*_ and the phase minimum *ϕ*_*min*_. (**b1**) The impedance *Z* is characterized by additional attributes: the anti-resonant frequency *f*_*ares*_, the impedance local minimum *Z*_*min*_ and *Q*_0_ = *Z*_*max*_ − *Z*_0_. The resonance amplitude is defined as *Q*_*Z*_ = *Z*_*max*_ − *Z*_*min*_ and it coincides with the definition in panel a (in the absence of antiresonance *Z*_*min*_ = *Z*_0_). (**b2**) The phase *ϕ* is characterized by two additional attributes: the phase local maximum *ϕ*_*max*_ and the zero-crossing phase *f*_*aphas*_ on the descending portion of *ϕ*.

#### 2.5.2 Impedance and phase profiles for 3D linear systems: emergence of antiresonance and antiphasonance

For certain parameter regimes (discussed later in this paper) the voltage response for 3D linear systems is more complex than for 2D linear ones [2, 18]. The impedance profile may exhibit a trough *Z*_*min*_ at an (anti-resonant) input frequency *f* = *f*_*ares*_ in addition to the peak at *f* = *f*_*res*_ (Fig. 1-b1). The phase profile may have an additional zero-frequency cross at *f* = *f*_*aphas*_ (antiphasonance) in addition to the one at *f* = *f*_*phas*_ (Fig. 1-b1). For *f* = *f*_*aphas*_ and *f* = *f*_*phas*_ the input current and output voltage peak at the same time. The voltage response is delayed for *f* < *f*_*aphas*_ and *f* > *f*_*phas*_ and advanced for *f* satisfying *f*_*phas*_ < *f* < *f*_*aphas*_.

#### 2.5.3 Attributes of the impedance and phase-profiles

The investigation of the role of resonant and amplifying currents requires the tracking of changes in both the impedance and phase profiles as a certain model parameter changes. It is helpful to characterize these graphs by using a small number of attributes (Fig. 1) that capture the salient properties of their shapes. Some of these attributes have been defined earlier: *f*_*res*_*, f*_*phas*_, *f*_*ares*_, *f*_*aphas*_, *Z*_*max*_ and *Z*_*min*_. When *f*_*ares*_ *=* 0, Z_*min*_ = *Z*_0_ (= *Z*(0)).

In addition, we will use the resonance amplitude *Q*_*Z*_ = *Z*_*max*_ − *Z*_*min*_ and the antiresonance amplitude *Q*_0_ = *Z*_*max*_ − *Z*_0_. In the absence of antiresonance, *Q*_*Z*_ = *Q*_0_. The half-width frequency band Λ_1/2_ is the length of the frequency interval in between *f*_*res*_ and the input frequency value at which *Z*(*f*) = *Z*_*max*_/2, and measures the system’s selectivity to input frequencies in the resonant frequency band. We do not explicitly analyze this attribute in this paper.

## 3 Results

The investigation of the mechanisms of generation of resonance, phasonance, antiresonance and antiphasonance consists primarily of understanding how and under what conditions *f*_*res*_), *f*_*phas*_*, f*_*ares*_) and *f*_*aphas*_ transition from zero to positive values respectively. Alternatively, resonance requires *Q*_*Z*_ > 0 and *Q*_0_ < *Q*_*Z*_ (for systems exhibiting resonance but not antiresonance *Q*_*Z*_ = *Q*_0_).

If a neuron exhibits resonance, the mechanisms of amplification of the voltage response in the resonant frequency band consist primarily of understanding how the resonance amplitude *Q*_*Z*_ and the maximal impedance *Z*_*max*_ increase as certain model parameter increases. These two attributes convey different information about the voltage response in the resonant frequency band. This is particularly important for neurons, since *Q*_*Z*_ may increase (decrease), while *Z*_*max*_ decreases (increases), thus bringing the voltage further away from (closer to) the threshold for spike generation in the resonant frequency band.

### 3.1 Mechanisms of generation of resonance and phasonance in 2D linear systems revisited

#### 3.1.1 Biophysical mechanisms

In [3,41] we identified the basic mechanisms of generation of resonance for the two-dimensional linear system (4)-(5) with *g*_2_ = 0 (henceforth, the 2D system). Additionally, we carried a thorough analysis of the properties of the attributes of the voltage response and their dependence on the linearized conductances, *g*_*L*_ and *g*_1_, and the time constant 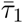. One lesson from these studies is that resonance and phasonance involve the complex interaction not only between the linearized conductances, but also the effective time scales. We revisit these results here and we refer the reader to [3, 41, 42] for a more detailed discussion.

Resonance in 2D linear systems can be created by three fundamental mechanisms involving: (i) an increase in *g*_1_, (ii) an increase in *g*_*L*_, and (iii) an increase in 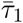. Relevant aspects of the dependences of the main attributes of the voltage response (*f*_*res*_, *f*_*phas*_, *Z*_*max*_ and *Z*_0_) with these parameters are summarized in Fig. 2.

**Figure 2:**
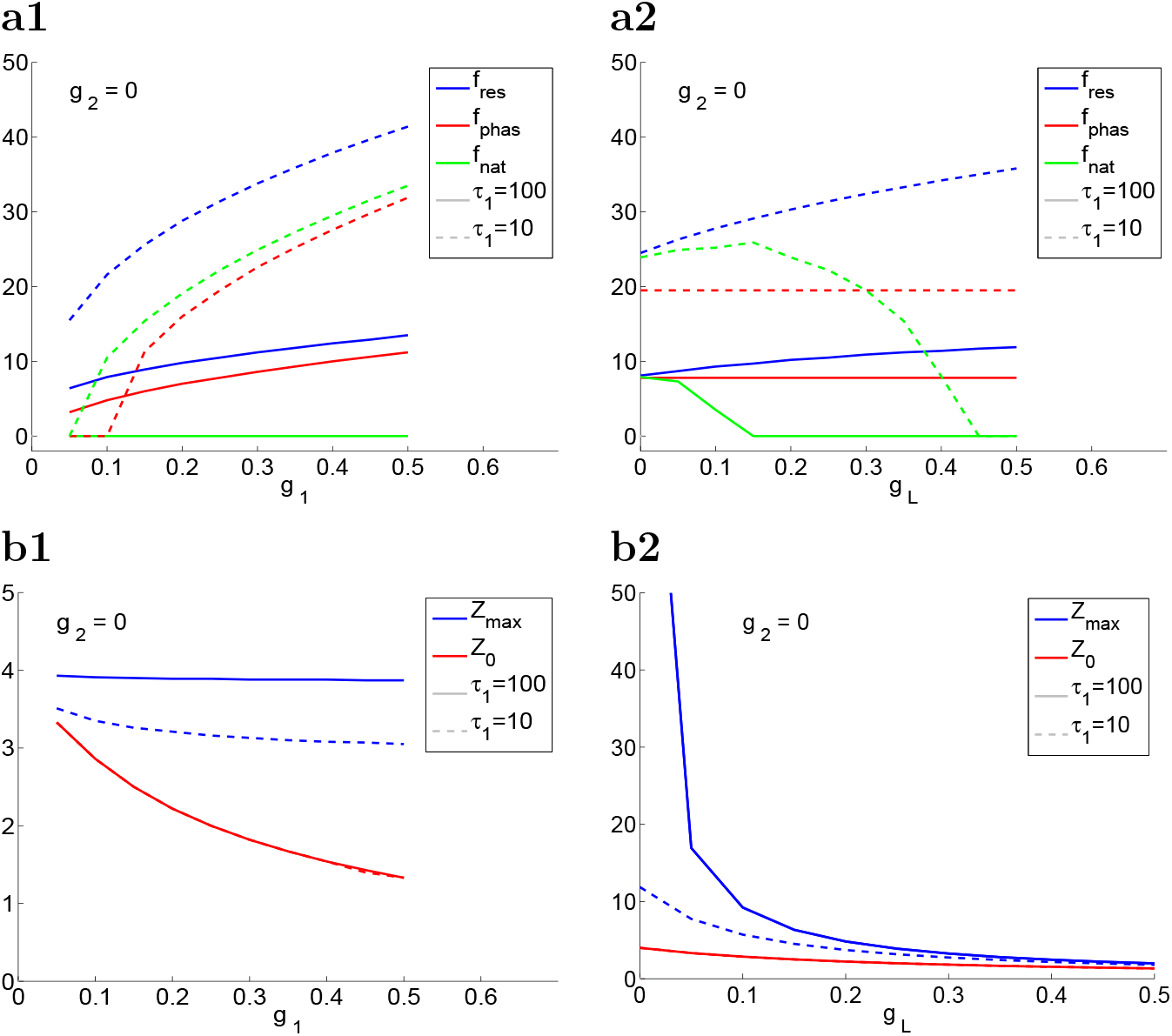
**Attributes of the voltage response for the 2D system (4)-(6) with** *g*_2_ = 0 **for representative parameter values.** (**a**) *f*_*res*_*, f*_*phas*_ and *f*_*nat*_ as a function of *g*_1_ for *g*_*L*_ = 0.25 (a1) and as a function of *g*_*L*_ for *g*_1_ = 0.25 (a2). (**b**) *Z*_*max*_ and *Z*_0_ as a function of *g*_1_ for *g*_*L*_ = 0.25 (b1) and as a function of *g*_*L*_ for g_1_ = 0.25 (b2). The superimposed red-solid and red-dashed curves indicate that *Z*_0_ is independent of *τ*_1_. We used the following parameter values *C* =1 and *A*_*in*_ = 1.

For *g*_1_ = 0, *f*_*res*_ = *f*_*phas*_ = *f*_*nat*_ = 0 and *Z*_*max*_ = *Z*_0_ (*Q*_*Z*_ = 0). As *g*_1_ increases above certain threshold value, resonance is generated by an unbalanced decrease in Z_*max*_ and *Z*_0_, which is more pronounced for *Z*_0_ (Fig. 2-b1). This threshold value for *g*_1_ depends on both *g*_*L*_ and 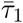. The 2D system may exhibit resonance, phasonance and intrinsic oscillations. If this happens, *f*_*res*_, *f*_*phas*_ and *f*_*nat*_ are increasing functions of *g*_1_ (Fig. 2-a1).

The ability of the 2D system to exhibit resonance for *g*_*L*_ = 0 depends on the values of both *g*_1_ and 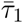. For the example in Fig. 2 the 2D system exhibits resonance for *g*_*L*_ = 0. While *f*_*res*_ is an increasing function of *g*_*L*_, *f*_*phas*_ is independent of *g*_*L*_ and *f*_*nat*_ is a decreasing function of *g*_*L*_ for a significantly large range of values of *g*_*L*_ (Fig. 2-a2). Resonance is amplified by a combined increase in both *Z*_*max*_ and *Z*_0_ as *g*_*L*_ decreases, which is significantly more pronounced for *Z*_*max*_ (Fig. 2-b2). While resonance is still present for large values of *g*_*L*_, *Q*_*Z*_ decreases very fast as *g*_*L*_ increases and approaches zero. This attenuation of the voltage response constitutes an effective annihilation of the resonance phenomenon.

The monotonic behavior of the resonance amplitude *Q*_*Z*_ is not necessarily associated with a specific monotonic behavior of *f*_*res*_ and *f*_*phas*_. In fact, *f*_*res*_ increases with increasing values of *g*_1_, but it first increases and then decreases with increasing values of *g*_*L*_ (Fig. 2-a).

The mechanism of generation of resonance by increasing values of 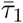 (and fixed values of both *g*_1_ and *g*_*L*_) involves an increase in *Z*_*max*_, while *Z*_0_ remains unchanged (Fig. 2-a2 and -b2). This amplification of the voltage response (compare the solid- and dashed-blue curves in Fig. 2-b) is accompanied by a decrease in *f*_*res*_*, f*_*phas*_ and *f*_*nat*_ (compare the corresponding solid and dashed curves in Fig. 2-a).

#### 3.1.2 Dynamic mechanisms

For the 2D system (*g*_2_ = 0), the properties of the voltage response to sinusoidal inputs result from the interplay of the cell’s intrinsic time scales and the input’s time scale. From (26)-(27) with *g*_2_ = 0, the former is determined by the effective time constants 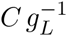 and 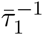. The latter is determined by the input frequency *f*. The time scale separation between *v* and *w*_1_ is given by 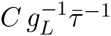. The parameter *g*_1_ (or 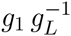) controls the strength of the negative feedback.

For *f* → 0, the cell response is instantaneous regardless of the values of the time constants. The voltage *v* is slaved to the current input. Therefore, the voltage maximum value is equal to the cell’s resistance (*Z*(0) = *Z*_0_ = (*g*_*L*_ + *g*_1_)^−1^) and the voltage and the input peak at the same time, so *ϕ*(0) = 0 (e.g., Figs. 3-a and -b). For very large values of *f*, on the other extreme, the voltage response is very slow as compared to the input’s speed. The voltage reaches very small extreme values and peaks with a delay approaching π/2. In the limit of *f* → ∞, regardless of the values of the time constants, the voltage response is zero (Figs. 3-a and -b).

**Figure 3:**
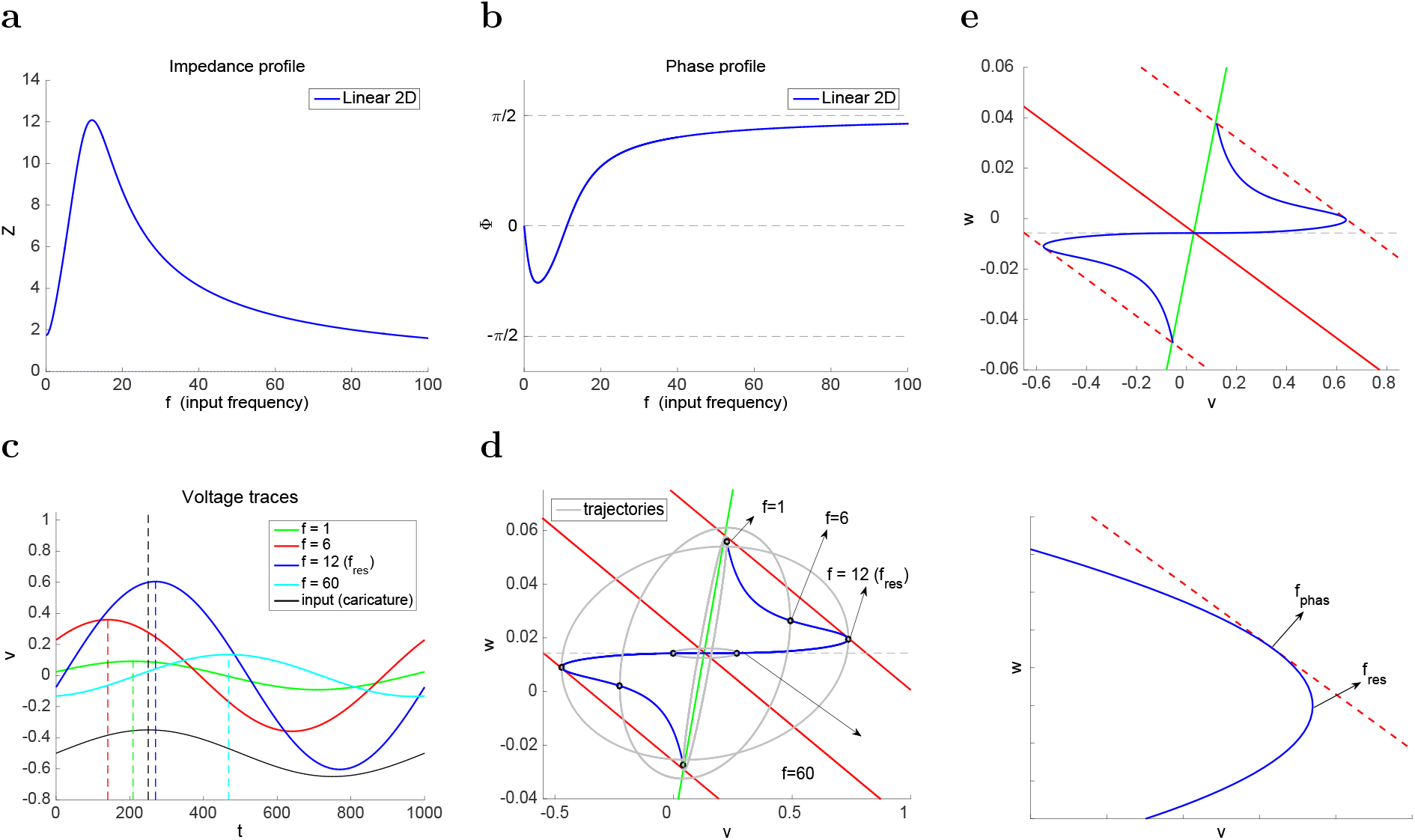
**Resonance and phasonance in 2D linear models.** **(a)** Impedance profile (*f*_*res*_ = 12). **(b)** Phase profile (*f*_*phas*_ = 11.1). The natural frequency is *f*_*nat*_ = 10. **(c)** Voltage traces for representative input frequency values. All curves are rescaled to have the same period (1000 ms). A caricature of the sinusoidal input (black curve) is included for reference below the curves. The dashed vertical lines note the peaks of the corresponding functions. The zero-phase response is indicated by the dashed-black vertical line (peak of the sinusoidal input). **(d)** Envelope-plane diagram. The red and green lines are the *v*- and *w*-nullclines respectively. The *v*-nullcline (solid-red) moves cyclically in between the displaced *v*-nullclines for ±*A*_*in*_ (dashed-red) following the sinusoidal input. The gray curves are the response limit cycle response trajectories corresponding to the voltage traces in panel c. The upper and lower envelope curves (solid blue curves) join the maximum (upper) and minimum (lower) points respectively on the response limit cycle response trajectories as *f* increases from *f* = 0 (intersection between the dashed-red and green lines) to *f* → ∞ (intersection between the solid-red and green lines determining the fixed-point). Because of linearity, the upper and lower envelope-curves are symmetric with respect to the fixed-point. **(e)** “Clean” envelope-plane diagram. The *v*-coordinates of the envelope-curve are *Z A*_*in*_. The maximum value of *v* on the envelope curve corresponds to *f* = *f*_*res*_. The point on the envelope curve tangent to the upper dashed-red line corresponds to *f* = *f*_*phas*_. The bottom panel is a magnification of the top one.

Resonance occurs if for an intermediate frequency range *Z*(*f*) > *Z*_0_, while phasonance requires the voltage response to be able to peak before the sinusoidal input (advanced response) for a range of input frequencies [41]. In both cases, the cell’s response is optimal at *f* = *f*_*res*_ and *f* = *f*_*phas*_, respectively, in the sense that it is neither too fast nor too slow to reach a maximum value and to peak in-phase with the input respectively. This requires appropriate combinations of the dimensionless parameters 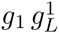 and 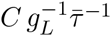 such that the former is large enough and the latter is small enough.

#### 3.1.3 The dynamic phase-plane

All this is better captured by the rescaled 2D model (30)-(31) with *κ* = 0 (henceforth the 2D model) and the dynamic phase-plane approach we used in [41] (see also [42] for an extension to 2D nonlinear systems). Fig. 3-c shows the superimposed voltage traces for representative input frequencies *f* (see panels a and b) during one cycle for the linear 2D model with a sinusoidal input of the form (3) for a representative value of *A*_*in*_. The black curve is a caricature of the sinusoidal input whose peak time serves as the phase reference (*ϕ* = 0).

Fig. 3-d shows the projection of the phase-space (for *v, w* and *t*) onto the *v-w* plane (henceforth the phase-plane) with the superimposed limit cycle response (LCR) trajectories corresponding to the voltage response curves for the input frequencies shown in Fig. 3-c. For clarity, we show the envelope-plane diagram without the LCR trajectories in Fig. 3-e.

We view the nullcline for the autonomous 2D model as the baseline for the analysis. As *t* progresses, this *v*-nullcline moves cyclically, parallel to itself in between the two red-dashed lines corresponding to *I*_*in*,1_ = *A*_*in*_ (top) and *I*_*in*,1_ = −*A*_*in*_ (bottom). Clearly, the distance between the moving *v*-nullcline and the baseline *v*-nullcline is equal to *A*_*in*_.

#### 3.1.4 Envelope-plane diagrams and envelope curves

The black dots in the LCR trajectories in Fig. 3-d note the points for which they reach their maximum *v*-values. As *f* increases these points span the blue curves, which we refer to as the upper and lower *envelope-curves* [41]. We refer to the diagrams such as these in Fig. 3-e, containing the *v*-nullclines (solid and dashed), the *w*-nullcline and the envelope-curves, as the *envelope-plane diagrams*.

Envelope-plane diagrams contain geometric and dynamic information about a system’s frequency response to oscillatory inputs, and are the frequency analogous to phase-plane diagrams [41]. Trajectories in the envelope-plane diagrams (upper and lower envelopes) are curves parameterized by the input frequency as trajectories in the phase-planes are curves parametrized by time. (Neither *f* nor *t* are explicit in these diagrams.) The cusp (“horizontal peak”) in the envelope curve corresponds to the peak in the impedance profile, and hence it corresponds to *f* = *f*_*res*_. The point on the envelope-plane curve tangent to the upper dashed-red line corresponds to *f* = *f*_*phas*_ since this tangency indicates that both the limit cycle trajectory and the v-nullcline reach their maximum at the same time.

The shape of the limit cycle trajectories depend on the interaction between the input frequency *f* and the underlying vector field [41]. We use this framework to explain the representative cases discussed above for the 2D linear system (30)-(31) (with *w*_1_ substituted by *w* and *κ* = 0)

#### 3.1.5 Quasi-one dimensional dynamics for low values of the input frequency *f*

For values of *f* ≪ 1 both equations are very fast (*dv/dt* → ∞ and *dw/dt* → ∞), and therefore the RLC trajectories track the motion of the fixed-point almost instantaneously (in a quasi-steady-state fashion). In the limit *f* → 0, the RLC trajectory moves cyclically along the *w*-nullcline in between the fixed-points generated by the dashed-red lines (intersection between these lines and the *w*-nullcline). The *v*-coordinate of this fixed-point is equal to (1 + *α*)^−1^ *A*_*in*_. For slightly higher values of *f*, such as *f* = 1, the RLC trajectories become elliptic-like around the *w*-nulcline, reflecting the emergence of a horizontal (off *w*-nullcline) direction of motion.

#### 3.1.6 Quasi-one dimensional dynamics for high values of the input frequency *f*

For large enough values of *f* ≫ 1, both equations are very slow (*dv/dt* → 0 and *dw/dt* → 0), and therefore the RLC trajectory evolves with a very low speed relative to the motion of the *v*-nullcline. Therefore, the *v*-nullcline “comes back” while the RLC trajectory has only reached a very small distance from the fixed-point and is forced to reverse direction. As a result, the amplitude of the RLC trajectory is small as compared to other values of *f* (e.g., limit cycle trajectory for *f* = 60). In the limit *f* → ∞, the amplitude of the RLC trajectory is zero (the limit cycle shrinks to the origin).

#### 3.1.7 Two-dimensional dynamics for input frequencies in the resonant frequency band

For intermediate values of *f* there is a transition in the shapes of the RLC trajectories between these two limit cases. Specifically, they first widen as their major axes rotate clockwise, and then they shrink while their major axis remains almost horizontal. Geometrically, the system exhibits resonance if for at least one of these RLC trajectories it is satisfied that *v*_*max*_(*f*) > *v*_*max*_ (0) as in Fig. 3-d for *f* = 6 and *f* = *f*_*res*_ = 12. The latter corresponds to the peak of the envelope-plane curve (Fig. 3-e). The system exhibits phase-resonance if both the *v*-nullcline and a RLC trajectory reach the upper dashed-red line at exactly the same time, thus generating a tangency between the envelope-plane curve and the dashed-red *v*-nullcline (Fig. 3-e). For the parameters in Fig. 3-d, the points on the envelope-curve corresponding to *f*_*res*_ and *f*_*phas*_ are very close (Fig. 3-e, bottom), almost indistinguishable.

For the limiting cases (*f* = 0 and *f* → ∞), the RLC trajectories are quasi one-dimensional. For intermediate values of *f* the RLC trajectories are neither too fast nor too slow, and then, while they are “left behind” by the moving *v*-nullcline, they can take advantage of the two-dimensional vector field without being constrained to move in quasi-one-dimensional directions. For the appropriate parameter values it is this degree of freedom that allows RLC trajectories to reach values of *v*_*max*_(*f*) larger than *v*_*max*_(0), and so to exhibit resonance.

### 3.2 Resonance and phasonance in 3D linear systems: limiting and special quasi-2D cases

The results for the 2D system described above can be used to explain the dynamics of the 3D linear system in three special cases for *τ*_2_: (i) *τ*_2_ → 0 (*w*_2_ has instantaneous dynamics), (ii) *τ*_2_ → ∞ (*w*_2_ has very slow dynamics), and (iii) *τ*_2_ = *τ*_1_ (*w*_1_ and *w*_2_ have identical dynamics). We assume that *τ*_1_ is away from *τ*_2_ in the first two cases.

In the first case, *w*_2_ is slaved to *v* (*w*_2_ ~ *v*) and, therefore, *g*_2_ can be absorbed into *g*_*L*_. A resonant (amplifying) gating variable will effectively increase (decrease) the value of *g*_*L*_, and therefore attenuate (amplify) the voltage response and increase (decrease) the resonant frequency. The envelope-curve lives in the *w*_1_ = *w*_2_ plane (Fig. 4-b).

**Figure 4:**
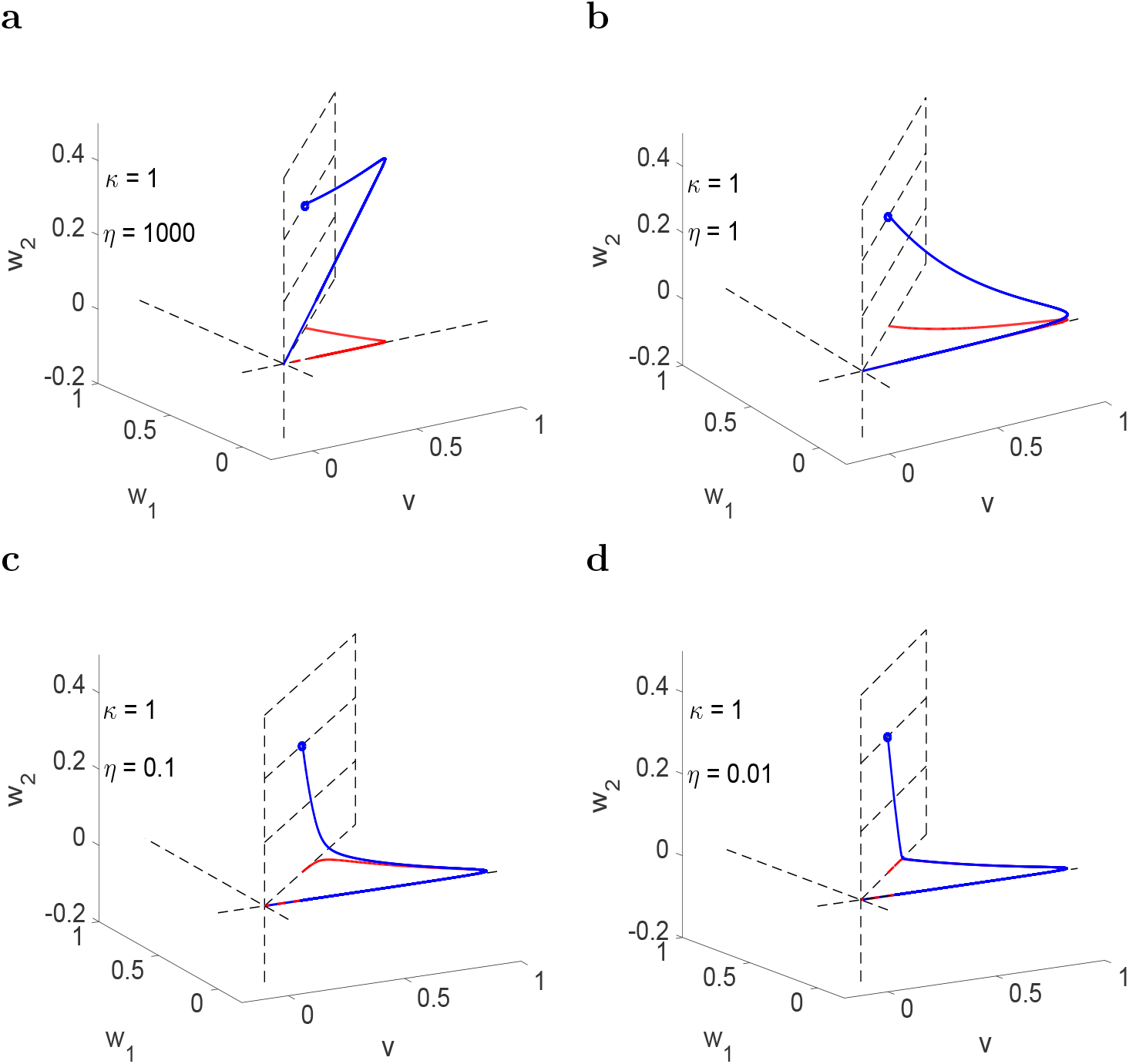
**Envelope-space diagrams for the linear 3D model in the quasi-2D regimes.** The (upper) envelope curves (solid blue) join the maximum points of the response limit cycle response trajectories as the input frequency *f* increases from *f* = 0 (blue dot on the *w*_1_-nullsurface) to *f* → ∞ (at the origin). The red curves are the projection of the envelope-curves onto the *v-w*_1_ (horizontal) plane. The lower envelope curves (not shown) are symmetric to the upper ones with respect to the origin (fixed-point of the autonomous system). We used the following parameter values: *α* = 1, *ϵ* = 0.05 and *κ* = −1.2. The dynamics become increasingly 2D as *η* decreases as indicated by the larger portions of the envelope-curves (blue) that are almost superimposed to their projections (red) onto the *v-w*_1_ planes. **(a)** *η* = 1000. The envelope curve “’lives” in the *v* = *w*_2_ plane. **(b)** *η* =1. The envelope curve “’lives” in the *w*_1_ = *w*_2_ plane. **(c)** *η* = 0.1. The envelope curve “lives” in the *v-w*_1_ plane, except for the zero frequency band. **(d)** *η* = 0. 01. The envelope curve “lives” in the *v-w*_1_ plane, except for the zero frequency band.

In the second case *η* → 0. Provided that *f* is away from zero, *w*_2_ has no dynamics (*dw*_2_/*dt* → 0). From the frequency-dependent properties point of view, system (4)-(6) behaves effectively as the 2D system described in Section 3.1 whose resistance is given by

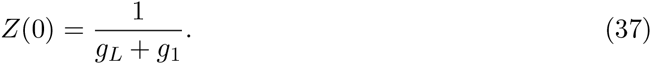

However, the 2D reduction is no longer valid when *f* < *η*, and therefore, a frequency boundary layer is created in the vicinity of *f* = 0 where the dynamics are quasi-1D and *Z*(0) is given by

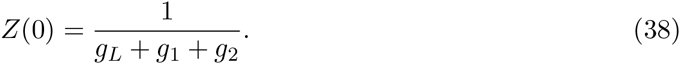

In other words, as *f* decreases *Z*(*f*) → (*g*_*L*_ + *g*_1_)^−1^, but *Z*(0) = (*g*_*L*_ + *g*_1_ + *g*_2_)^−1^. As we will discuss later in the paper, the abrupt transition required by the continuity of the solution to match these two values as *f* → 0 is the main component of the generation of antiresonance and antiphasonance. Away from this frequency band, the envelope-curve lives in the (horizontal) *v-w*_1_ plane (Figs. 4-c and -d).

In the third case, after transients have disappeared *w*_2_ = *w*_1_, then the two terms *g*_1_ *w*_1_ + *g*_2_ *w*_2_ in eq. (4) can be combined into one with *g*_1_ replaced by *g*_1_ + *g*_2_. A resonant (amplifying) gating variable *x*_2_ will effectively increase (decrease) the resonant effect of *w*_1_ and therefore increase (decrease) the resonant frequency and increase (decrease) the resonance amplitude. The envelope-curve lives in the *v* = *w*_2_ plane (Fig. 4-a).

The resulting modulatory effects on the dynamics of he 2D system caused by values of *g*_2_ ≠ 0 are expected to persist for values of *τ*_2_ away from these special cases, but close enough to them, where the dynamics is effectively quasi-2D. In the next Section we examine how changes in *τ*_2_ affects the behavior of the the voltage response as its qualitative behavior transitions through the cases discussed above.

### 3.3 Modulation and annihilation of existing resonances and emergence of antiresonance: basic biophysical mechanisms

Here we investigate the consequences of negative (*g*_2_ > 0) and positive (*g*_2_ < 0) feedback effects on the voltage response for the 3D system. We chose a 2D baseline system (*g*_2_ = 0) in a parameter regime (*C* = 1, *g*_*L*_ = 0.25, *g*_1_ = 0.25 and *τ*_1_ = 100) for which it exhibits both resonance and phasonance (blue curves in Fig. 5-a). We examine how changes in the linearized conductance *g*_2_ affect the voltage response for *g*_2_ = 0 for representative values of the time constant *τ*_2_ that increases from the top to the bottom panels. These values have been chosen to be below and comparable to *τ*_1_. We divided our study into two cases: (i) interaction between two negative feedback effects (*g*_2_ > 0, Fig. 5), and (ii) interaction between a negative and a positive feedback effects (*g*_2_ < 0, Fig. 6). In both the cooperative (*g*_2_ < 0) and competitive (*g*_2_ < 0) feedback cases the voltage response is affected in qualitatively different ways depending on the values of *τ*_2_ (relative speed of the two processes). However, these differences are more pronounced for the latter.

**Figure 5:**
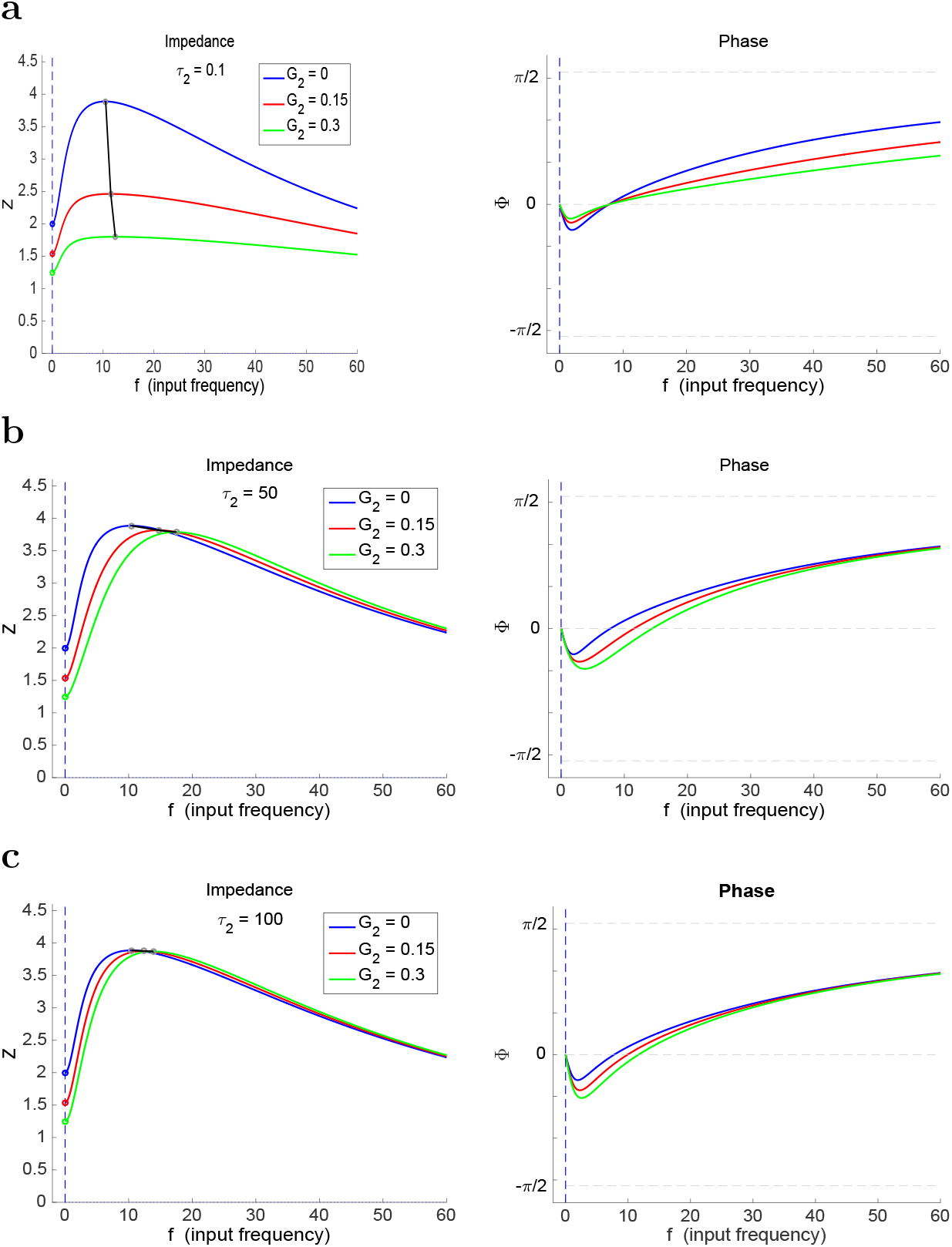
**Impedance and phase profiles for system (4)-(6) with** *g*_2_ > 0 **(interaction between two resonant feedback effects) for representative sets of parameter values.** The gray lines in the left panels connect the peaks of the impedance profiles. We used the following parameter values *C* = 1, *g*_*L*_ = 0.25, *g*_1_ = 0.25, *τ*_1_ = 100 and *A*_*in*_ = 1.

#### 3.3.1 Interaction between two negative feedback effects: modulation of the voltage response

The interaction between two negative feedback effects (*g*_2_ > 0) modulates the voltage response without generating antiresonances (Fig. 5). The “direction” and properties of this modulation depends on the value of *τ*_2_. When *w*_2_ is fast (*τ*_2_ = 0.1, Fig. 5-a), an increase in *g*_2_ causes a strong attenuation of the voltage response accompanied by slight changes in *f*_*res*_ and *f*_*phas*_. This is expected from our previous discussion in Section 3.2. This attenuation involves a strong decrease in *Z*_*max*_ and a smaller decrease in *Z*_0_.

In contrast, for larger values of *τ*_2_, there is an amplification of the voltage response as *g*_2_ increases that is caused by an increase in *Q*_*z*_ due to a smaller decrease in *Z*_*max*_ than in *Z*_0_ (Figs. 5-b and -c). This is accompanied by increases in *f*_*res*_ and *f*_*phas*_. These effects are more pronounced for *τ*_2_ = 50 (< *τ*_1_) (Fig. 5-b) than for *τ*_2_ = *τ*_1_ = 100 (Fig. 5-b). As *τ*_2_ increases further, *Z*_*max*_ approaches a constant value (not shown) and so does *Q*_*Z*_ since *Z*_0_ is independent of *τ*_2_. As expected, the impedance and phase profiles for different values of *g*_2_ almost coincide for larger values of *f* except in the lowest frequency band whose size decreases with increasing values of *τ*_2_ (compare Figs. 5-b and -c).

From (38), an increase (decrease) in *g*_2_ causes a decrease (increase) in *Z*_0_. In a separate set of simulations we have considered combined changes in the values of *g*_1_ and *g*_2_ such that *g*_1_ + *g*_2_ remains constant (*g*_1_ + *g*_2_ = 0.25) and, therefore, *Z*_0_ also remains constant. The results (not shown) are qualitatively similar to these presented here (Figs. 5 and 6). One difference is that the impedance and phase profiles coincide along these level sets for *τ*_2_ = *τ*_1_ = 100 and there is a small amplification of the voltage response as with increasing values of *g*_2_ as *τ*_2_ increases above *τ*_2_ = 100. This is not surprising since *g*_1_ decreases as *g*_2_ increases and the activation of *w*_2_ is delayed as compared to *w*_1_. This amplification of the voltage response is accompanied with small decreases in both *f*_*res*_ and *f*_*phas*_.

**Figure 6:**
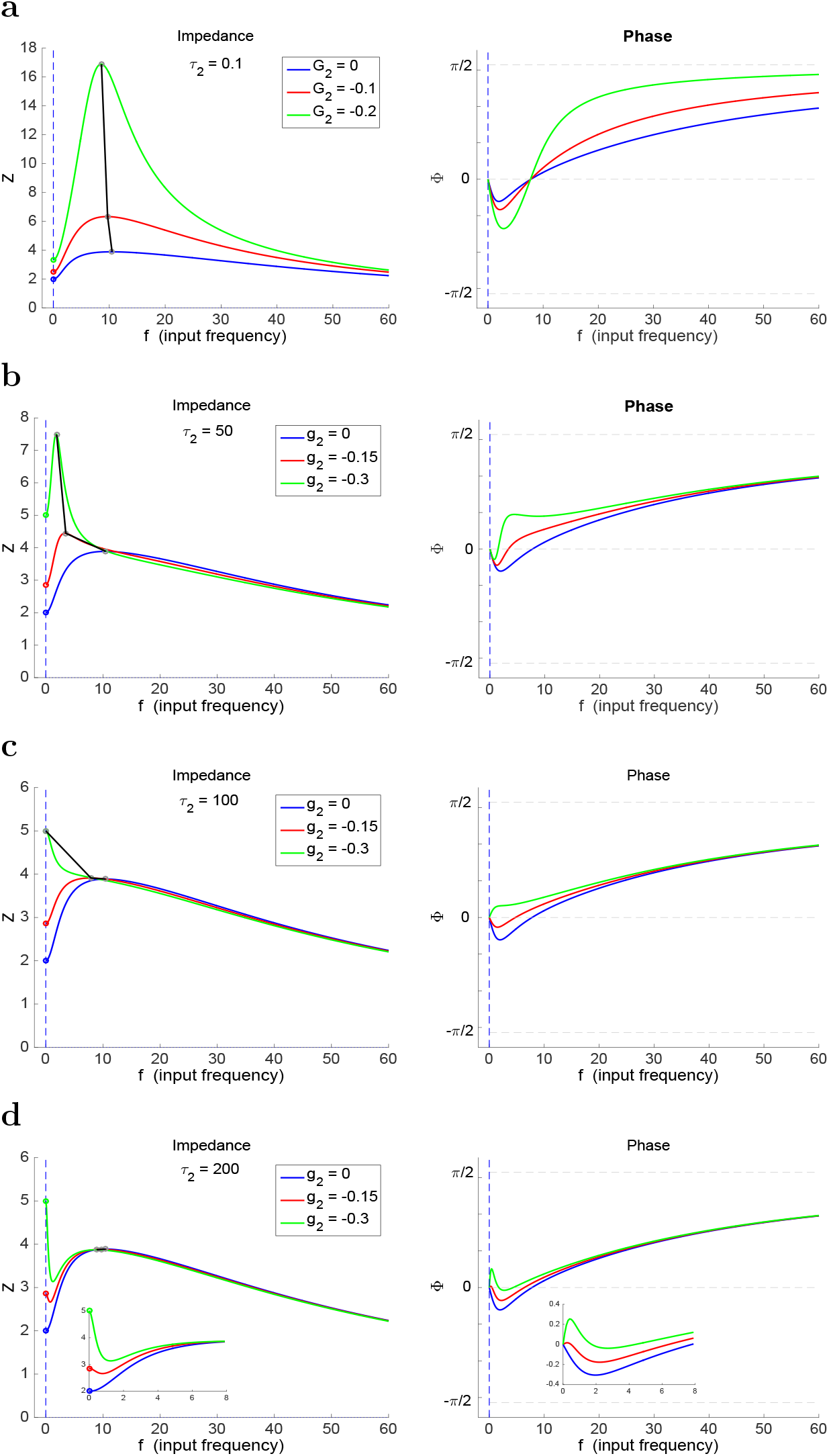
**Impedance and phase profiles for system (4)-(6) with** *g*_2_ < 0 **(interaction between a negative and a positive feedback effects) for representative sets of parameter values.** The gray lines in the left panels connect the peaks of the impedance profiles. We used the following parameter values *C* = 1, *g*_*L*_ = 0.25, *g*_1_ = 0.25, *τ*_1_ = 100 and *A*_*in*_ = 1.

#### 3.3.2 Interaction between a slow negative and a relatively fast positive feedback effects: amplification of the voltage response

For low values of *τ*_2_, an increase in |*g*_2_| produces a strong amplification of the voltage response (Fig. 6-a) and a slight decrease in *f*_*res*_ and *f*_*phas*_. For larger values of *τ*_2_ < *τ*_1_ the amplification of the voltage response as |*g*_2_| increases is less pronounced than for *τ*_2_ = 0.1 (Fig. 6-b) and both *f*_*res*_ and *f*_*phas*_ decrease. The weaker amplification of the voltage response results from a smaller increase in *Z*_*max*_ and a larger increase in *Z*_0_ as |*g*_2_| increases. This in turn results from the slower activation of the positive feedback. This trend continues as long as *τ*_2_ is below *τ*_1_.

#### 3.3.3 Interaction between negative and positive feedback effects with comparable slow dynamics: annihilation of resonance

As *τ*_2_ approaches *τ*_1_ both resonance and phasonance are annihilated as *g*_2_ decreases (|*g*_2_| increases) past a threshold value (Fig. 6-c). For the green curve in Fig. 6-c, the voltage response at non-zero frequencies is not fast enough to raise above *Z*_0_ and, therefore, *Q*_*Z*_ = 0.

Note that the slower activation of the positive feedback renders the voltage response almost unaffected by increasing values of |*g*_2_| for values of *f* outside a small vicinity of *f* = 0 whose size decreases with increasing values of *τ*_2_ (compare Figs. 6-b, -c and -d, left panels).

#### 3.3.4 Interaction between a slow and a slower negative feedback effects: generation of antiresonance and antiphasonance

As *τ*_2_ increases above *τ*_1_, both antiresonance and antiphasonance emerge (Fig. 6-d). As for *τ*_2_ = 100 (Fig. 6-c), for *τ*_2_ = 200 (Fig. 6-d) the slower activation of *w*_2_ renders the values of *Z*_*max*_ almost unaffected by increasing values of |*g*_2_|. However, the interaction between a faster negative feedback and a slower positive feedback causes *Z*(*f*) to decrease below *Z*_*max*_ before reaching this value. This antiresonance mechanism restitutes resonance. An analogous effect is observed in the phase profiles where antiphasonance emerges as |*g*_2_| increases for *τ*_2_ > *τ*_1_.

For values of *g*_1_ and *g*_2_ such that *g*_1_ + *g*_2_ remain constant (not shown), the picture does not qualitatively change for values of *τ*_2_ < *τ*_1_. For *τ*_2_ = *τ*_1_ = 100, the impedance and phase profiles coincide. antiresonance and antiphasonance still emerge for values of *τ*_2_ = 200, but both phenomena are much less pronounced that in the case described in the previous paragraphs.

### 3.4 The shaping of the attributes of the impedance and phase profiles by the complex interplay of *g*_2_ and 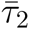

Here we extend the results from the previous sections to provide a global picture of how changes in *g*_2_ and *τ*_2_ affect the voltage response of the 3D linear system to sinusoidal inputs. Our results are presented in Figs. 7 to 11 for the same set of baseline parameter values as in Figs. 5 and 6. The colormap diagrams code for the values attributes of the impedance and phase profiles in the *g*_2_ - *τ*_2_ parameter space for fixed-values of the baseline parameters. The other graphs in each figure correspond to representative curves of for these attributes as a function of *τ*_2_ and *g*_2_. In these graphs we include the corresponding curves for *τ*_1_ = 10 (dashed) in addition to the (baseline) *τ*_1_ = 100 (solid).

**Figure 7:**
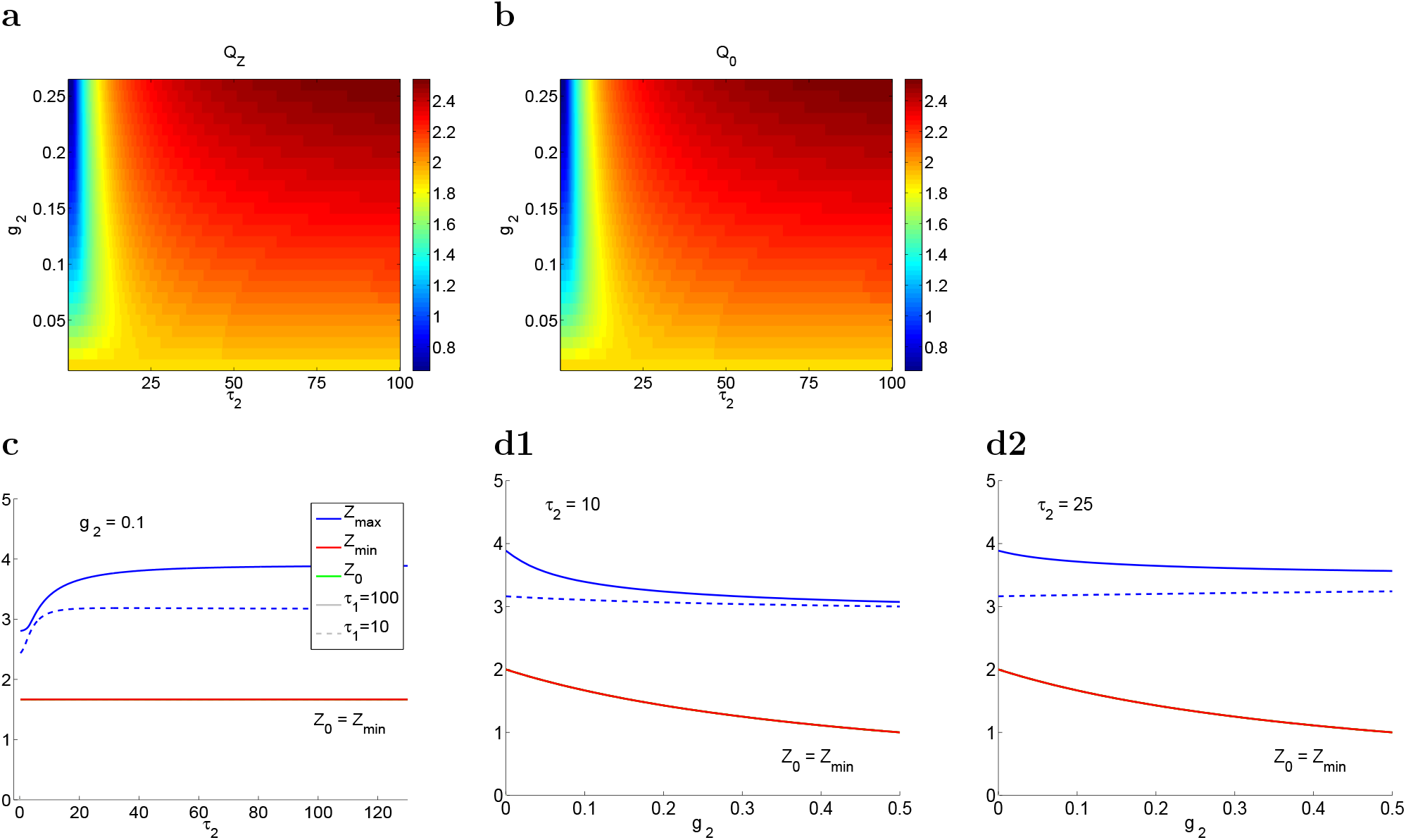
**Resonance amplitudes for system (4)-(6) with a resonant gating variable** *x*_2_ (*g*_2_ > 0) **for a representative set of parameter values.** **Top panels (a, b):** Colormap diagrams for *Q*_*Z*_ (a) and *Q*_0_ (b) in *Gg*_2_ - *τ*_2_ parameter space for *τ*_1_ = 100. **Bottom panels (c, d):** Representative curves of *Z*_*max*_, *Z*_0_ and *Z*_*min*_ as a function of *τ*_2_ (c) and *g*_2_ (d) for *τ*_1_ = 100 (solid) and *τ*_1_ = 10 (dashed). The superimposed red and green curves indicate that no antiresonances are present. The superimposed green-solid and green-dashed curves indicate that *Z*_0_ (and *Z*_*min*_) does not change with *τ*_2_. We used the following parameter values *C* = 1, *g*_*L*_ = 0.25, *g*_1_ = 0.25 and *A*_*in*_ = 1.

#### 3.4.1 Dependence of the modulating effects of the resonant gating variable *x*_2_ (*g*_2_ > 0) on *g*_2_ and *τ*_2_

For values of *g*_2_ > 0 the 3D system exhibits no antiresonances (Figs. 7-a and -b, showing that *Q*_*Z*_ and *Q*_0_ are identical). The impedance and phase profiles are qualitatively similar to these for the 2D system. The presence of an additional resonant gating variable modulates the voltage response and the resonance and phasonance phenomena, though not always in obvious ways.

As discussed above, the effect of changes in the values of *g*_2_ on the resonant properties of the 3D system depends on the values of *τ*_2_ (Fig. 7-a). For large enough values of *τ*_2_, increasing *g*_2_ causes an increase in *Q*_*Z*_, similarly to its dependence on *g*_1_ described above. This dependence is inverted for low values of *τ*_2_: in the presence of a fast negative feedback, increasing *g*_2_ causes an attenuation of the voltage response and a decrease in *Q*_*Z*_.

#### 3.4.2 The modulation mechanisms of the resonant gating variable *x*_2_ (*g*_2_ > 0) depend on *τ*_1_

We show this in Fig. 7-d. For large enough values of *τ*_1_ (solid curves), the increase in *Q*_*Z*_ as *g*_2_ increases results from a combined decrease in both *Z*_*max*_ and *Z*_0_, which is more pronounced for *Z*_0_ than for *Z*_*max*_. In contrast, for low enough values of *τ*_1_ (dashed lines), *Z*_*max*_ remains constant as *g*_2_ increases, while *Z*_0_ decreases.

Increasing the value of *τ*_2_ causes an amplification of the voltage response and an increase in *Q*_*Z*_, which depend on the value of *g*_2_. As *τ*_2_ increases, the negative feedback effect provided by *x*_2_ is slower, and therefore the voltage response can increase to higher values before this increase is opposed by *x*_2_. This amplification is more pronounced for larger values of *τ*_1_ (Fig. 7-c).

#### 3.4.3 Increasing values of *τ*_2_ generate bumps in the *f*_*res*_ and *f*_*phas*_ patterns

Both *f*_*res*_ and *f*_*phas*_ increase with increasing values of *g*_2_ (Figs. 8-a and -b) with *f*_*res*_ > *f*_*phas*_ (Fig. 8-d). Their values and rates of change depend on *τ*_2_. Following our discussion in Section 3.2, their dependence can be thought of a transition from the values corresponding to *τ*_2_ → 0 (2D system with *g*_*L*_ substituted by *g*_*L*_ + *g*_2_) to the values corresponding to *τ*_2_ → ∞ (2D system with *g*_2_ = 0). This transition is non-monotonic and involves an increase in both *f*_*res*_ and *f*_*phas*_ for low enough values of *τ*_2_ followed by a decrease as *τ*_2_ continues to increase (Fig. 8-d). These properties of these “bumps” depend not only on the value of *g*_2_ (Figs. 8-a and -b) but also on the value of the *τ*_1_: the smaller *τ*_1_ the stronger the bumps.

**Figure 8:**
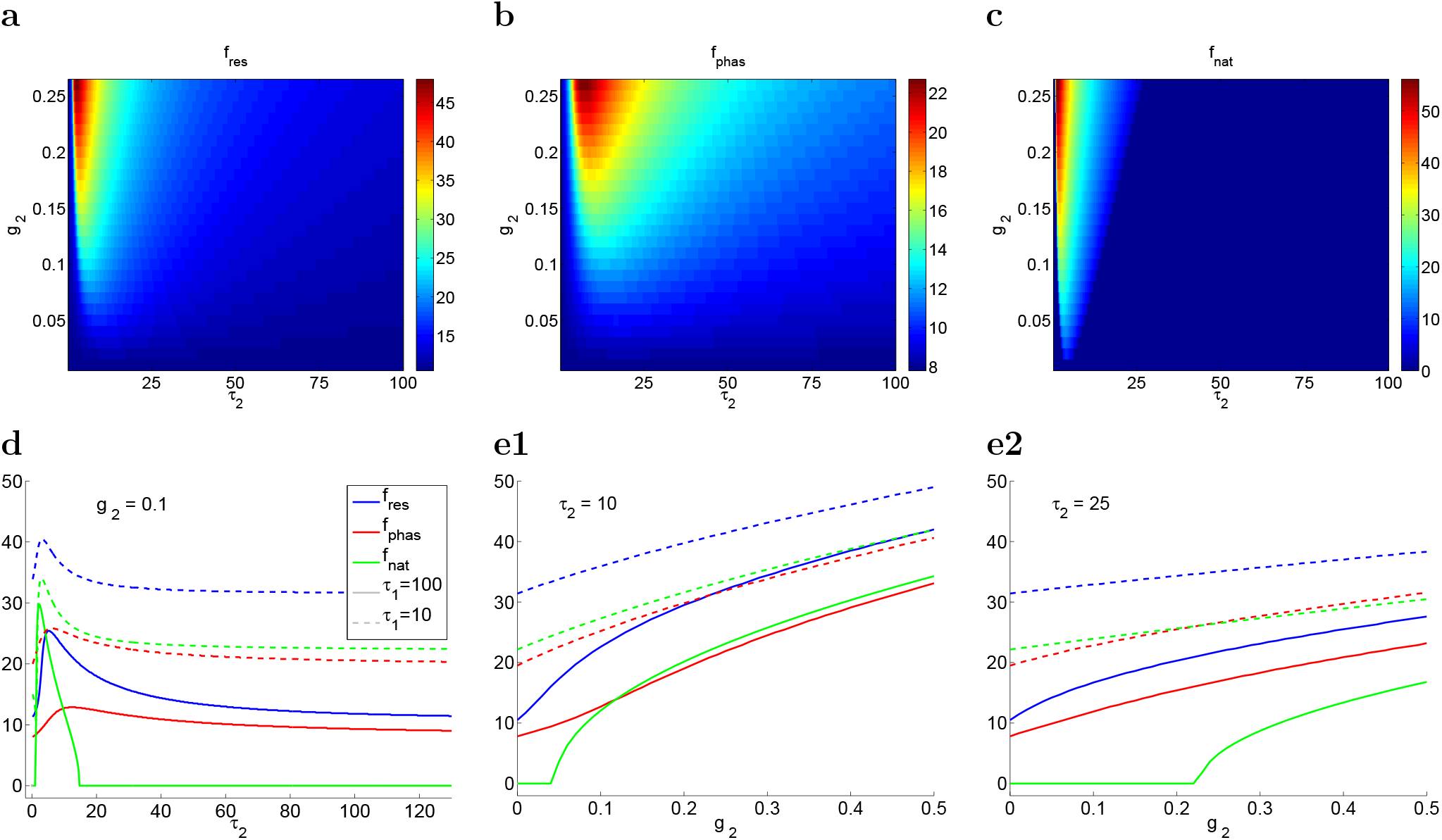
**Resonant, phasonant and natural frequencies for system (4)-(6) with a resonant gating variable** *x*_2_ (*g*_2_ *>* 0) **for a representative set of parameter values.** **Top panels (a, b, c):** Colormap diagrams for *f*_*res*_ (a), *f*_*phas*_ (b) and *f*_*nat*_ (c) in *g*_2_ - *τ*_2_ parameter space for *τ*_1_ = 100. **Bottom panels (d, e):** Representative curves of *f*_*res*_, *f*_*phas*_ and *f*_*nat*_ as a function of *τ*_2_ (d) and *g*_2_ (e) for *τ*_1_ = 100 (solid) and *τ*_1_ = 10 (dashed). We used the following parameter values *C* =1, *g*_*L*_ = 0.25, *g*_1_ = 0.25 and *A*_*in*_ = 1.

#### 3.4.4 Antiresonances are generated by the interplay of a slow resonant (*g*_1_ > 0) and a slower amplifying (*g*_2_ < 0)) gating variables

For values of *g*_2_ < 0 antiresonance emerge provided *τ*_2_ is large enough and *g*_2_ negative enough (large enough in absolute value). This is reflected in the different colormap diagrams for *Q*_*Z*_ and *Q*_0_ (Figs. 9-a and -b) in these parameter regimes. Figs. 9-d show the dependence of *Q*_*Z*_ and *Q*_0_ with *g*_2_ for representative values of 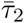 and 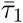. Since *Z*_0_ is independent of 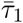 and 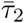, the solid and dashed red curves (*τ*_1_ = 100 and *τ*_1_ = 10, respectively) coincide in panels c and d. In addition, the red and green curves coincide in panels d1 (left and right, respectively).

**Figure 9:**
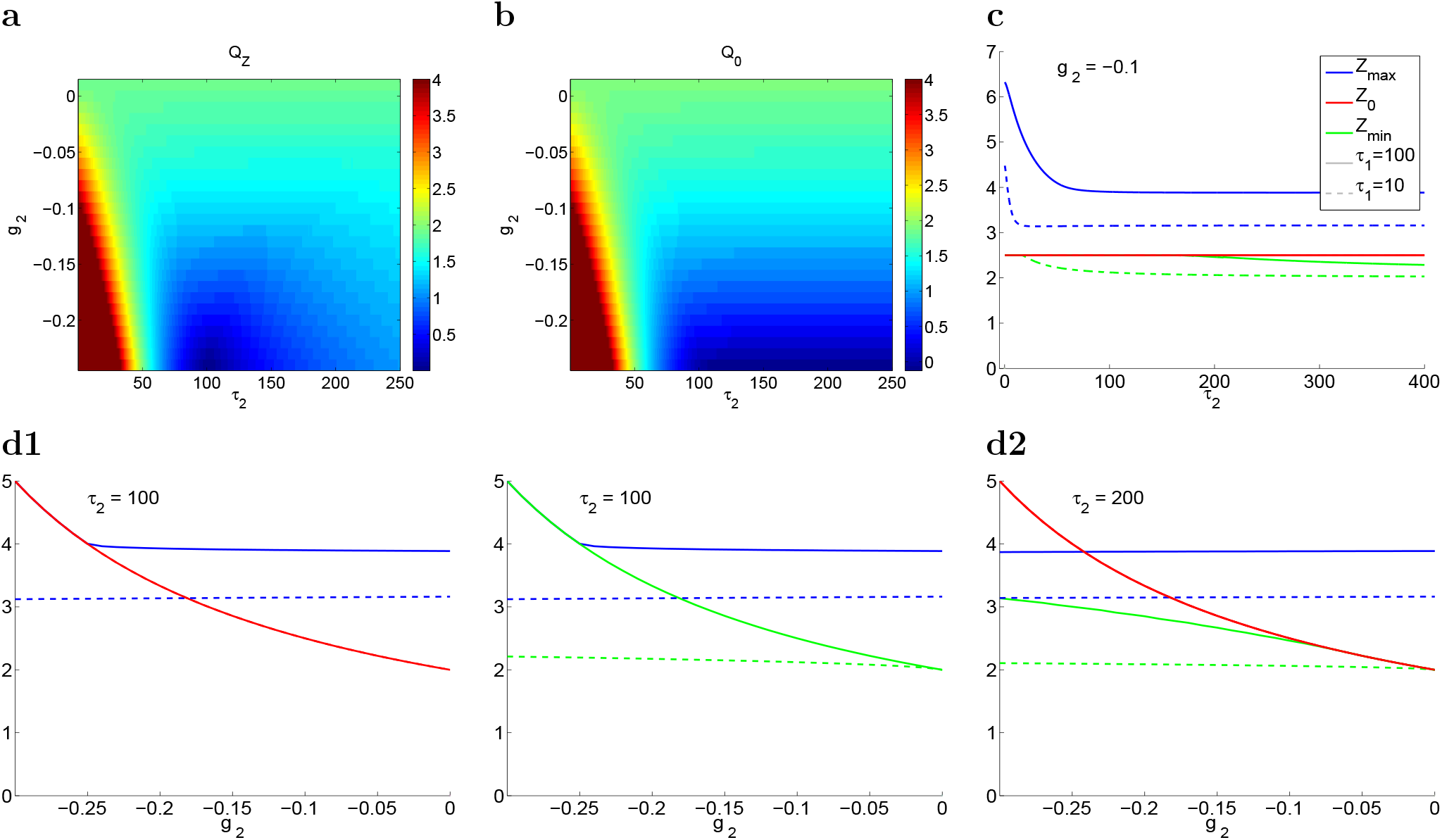
**Resonance amplitudes for system (4)-(6) with an amplifying gating variable** *x*_2_ (*g*_2_ < 0) **for a representative set of parameter values.** **Panels a, b:** Colormap diagrams for *Q*_*Z*_ (a) and *Q*_0_ (b) in *G*_2_ - *τ*_2_ parameter space for *τ*_1_ = 100. **Panels c, d:** Representative curves of *Z*_*max*_, *Z*_0_ and *Z*_*min*_ as a function of *τ*_2_ (c) and *h*_2_ (d) for *τ*_1_ = 100 (solid) and *τ*_1_ = 10 (dashed). The superimposed red-solid and red-dashed lines indicate that *Z*_0_ does not depend on *τ*_2_. We used the following parameter values *C* = 1, *G*_*L*_ = 0.25, *G*_1_ = 0.25 and *A*_*in*_ = 1.

For *τ*_1_ = 100 and *τ*_2_ = 100 (panels d1, solid lines) *Z*_*min*_ = *Z*_0_ (*Q*_*Z*_ = *Q*_0_) for all values of *g*_2_. In contrast, for *τ*_1_ = 100 and *τ*_2_ = 200 (panel d2, solid lines), the solid red (*Z*_0_) and green (*Z*_*min*_) curves separate as *g*_2_ decreases, indicating the generation of antiresonances. Note that *Q*_*Z*_ = *Z*_*max*_ − *Z*_0_ < 0 for the lowest values of *g*_2_ in panel d2 (see also 6-d). For *τ*_1_ = 10 (dashed curves), the antiresonances emerge for lower values of *τ*_2_ and higher values of *g*_2_ (panels d, dashed curves, red-dashed and -solid curves are superimposed). These results show that the trade-off between currents for the generation of resonance does not only involve the conductances, voltage-dependencies and time scales associated to the amplifying gating variable but also the properties of the resonant gating variable *x*_1_.

#### 3.4.5 *f*_*res*_ and *f*_*phas*_ decrease with decreasing values of *g*_2_ < 0, but have a non-monotonic (“anti-bump-like”) dependence with *τ*_2_

Fig. 10 shows that both *f*_*res*_ and *f*_*phas*_ decrease with decreasing values of *g*_2_ < 0 with rates that depend on the value of 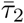 (Fig. 10-a and -b). Similarly to the case *g*_2_ > 0, the transition from 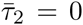 to 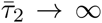 is non-monotonic and involves a decrease and subsequent increase in both *f*_*res*_ and *f*_*phas*_ in a relatively small range of values of 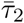. This range increases with decreasing values of 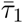 (compare the solid and dashed curves in Fig. 10-d for 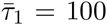 and 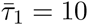 respectively). The changes in the values of *f*_*res*_ and *f*_*phas*_ within these ranges are more pronounced for 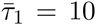 (dashed curves) than for *τ*_1_ = 100 (solid curves) respectively. The dependence of *f*_*res*_ and *f*_*phas*_ with *g*_2_ is monotonic (Fig. 10-e).

**Figure 10:**
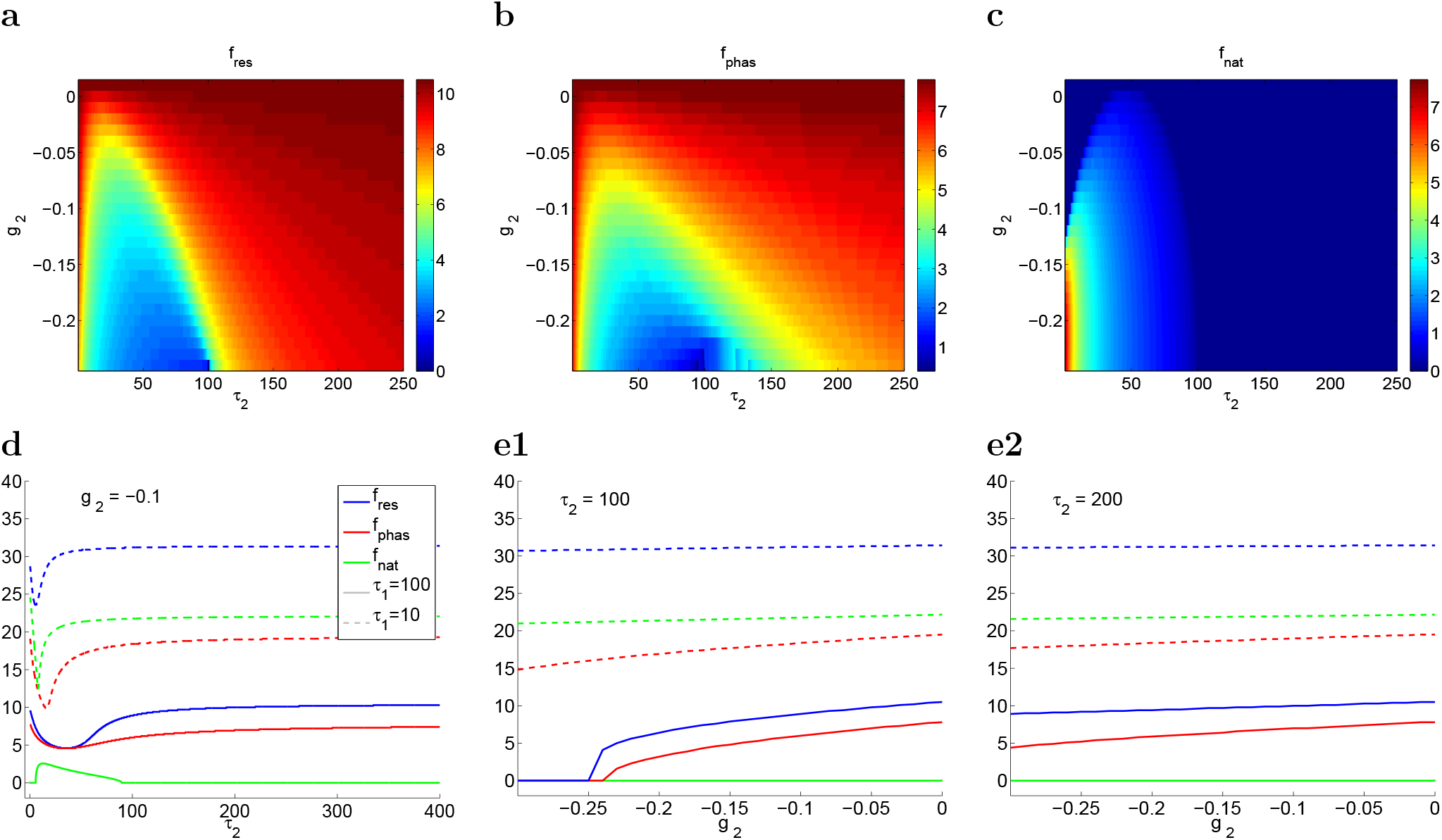
**Resonant, phasonant and natural frequencies for system (4)-(6) with an amplifying gating variable** *x*_2_ (*g*_2_ < 0) **for a representative set of parameter values.** **Top panels (a, b, c):** Colormap diagrams for *f*_*res*_ (a), *f*_*phas*_ (b) and *f*_*nat*_ (c) in *g*_2_ - *τ*_2_ parameter space for *τ*_1_ = 100. **Bottom panels (d, e):** Representative curves of *f*_*res*_, *f*_*phas*_ and *f*_*nat*_ as a function of *τ*_2_ (d) and *g*_2_ (e) for *τ*_1_ = 100 (solid) and *τ*_1_ = 10 (dashed). We used the following parameter values *C* =1, *g*_*L*_ = 0.25, *g*_1_ = 0.25 and *A*_*in*_ = 1.

#### 3.4.6 *f*_*ares*_ and *f*_*aphas*_ increase with decreasing values of *g*_2_ < 0, but have a non-monotonic (“bump-like”) dependence with *τ*_2_

Fig. 11 illustrates the dependence of *f*_*ares*_ and *f*_*aphas*_ with *g*_2_ and 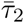 for representative parameter values. Comparison between Figs. 11-a and -b shows that antiresonance may exist in the absence of antiphasonance. Comparison between Figs. 11 and 10 shows that *f*_*ares*_ and *f*_*aphas*_ are always smaller than *f*_*res*_ and *f*_*phas*_. Similarly to *f*_*res*_ and *f*_*phas*_, *f*_*ares*_ and *f*_*aphas*_ are larger the smaller the values of 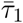. While typically *f*_*res*_ > *f*_*phas*_, for a given set of parameter values, there are parameter regimes where *f*_*ares*_ < *f*_*aphas*_ (Fig. 11-c, dashed curves). The dependence of *f*_*ares*_ and *f*_*aphas*_ with *τ*_2_ is not necessarily monotonic and may exhibits “bumps” where these quantities first increase and then decrease. The dependence of *f*_*ares*_ and *f*_*aphas*_ with *g*_2_ is monotonic, but *f*_*aphas*_ typically decrease faster than *f*_*ares*_ and these curves may intersect (Fig. 11-d).

**Figure 11:**
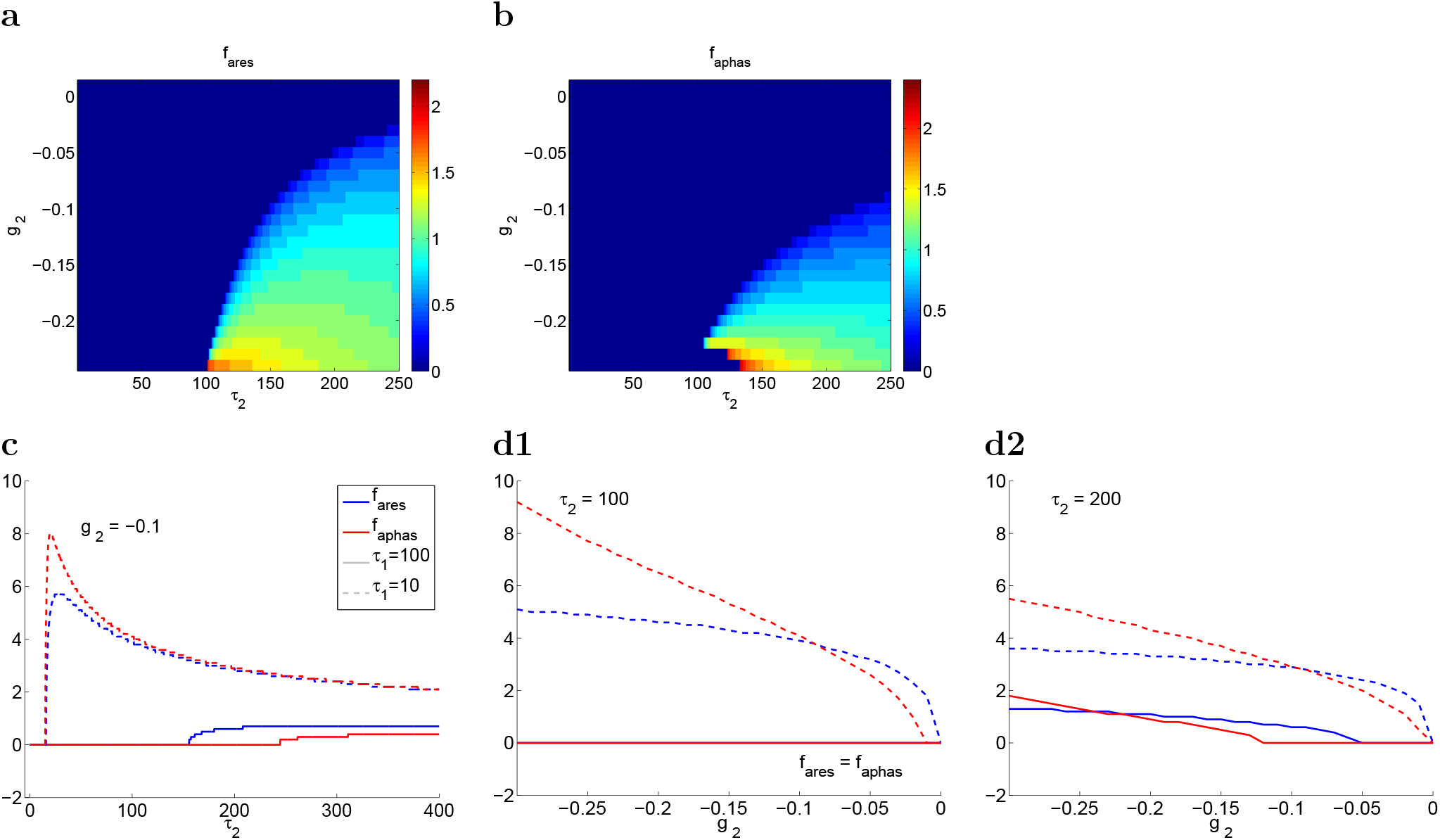
**Antiresonant and antiphasonant frequencies for system (4)-(6) with an amplifying gating variable** *x*_2_ (*G*_2_ < 0) **for a representative set of parameter values.** **Top panels (a, b):** Colormap diagrams for *f*_*ares*_ (a) and *f*_*aphas*_ (b) in *g*_2_ - *τ*_2_ parameter space for *τ*_1_ = 100. **Bottom panels (c, d):** Representative curves of *f*_*ares*_ and *f*_*aphas*_ as a function of *τ*_2_ (c) and *g*_2_ (d) for *τ*_1_ = 100 (solid) and *τ*_1_ = 10 (dashed). In panel d1, *f*_*res*_ = *f*_*aphas*_ = 0. We used the following parameter values *C* =1, *g*_*L*_ = 0.25, *g*_1_ = 0.25 and *A*_*in*_ = 1.

### 3.5 Dynamic mechanisms of generation of antiresonance and antiphasonance

Here we extend the analysis of 2D linear systems introduced in Section 3.1 to explain the mechanisms of generation of antiresonance and antiphasonance in the 3D linear system. We use the rescaled model (30)-(32) introduced in Section 2.3.2 (geometric rescaling). From our discussion in Section 3.4.4 and Fig. 6 (see also Section 3.2) antiresonance occurs when (i) *x*_1_ is resonant (*g*_1_ > 0), and (ii) *x*_2_ is amplifying (*g*_2_ < 0) and slower than *x*_1_ (*τ*_2_ < *τ*_1_). Assuming *g*_*L*_ > 0, this yield *α* > 0 and *κ* < 0. We will focus on the transition of *η* from *η* > 1 (*τ*_2_ > *τ*_1_) to *η* < 1 (*τ*_2_ < *τ*_1_) as we did in Fig. 4. We will use parameter values similar to the ones used in the previous sections: *α* = 1 and *ϵ* = 0.05.

As discussed above and shown in Fig. 6, one immediate consequence of the presence of the second amplifying gating variable (*κ* < 0) is an increase in the cell’s resistance, given by

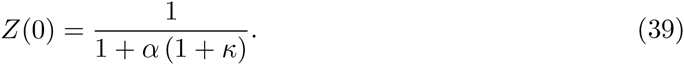

This has different consequences depending on the values of *η* and *κ*: resonance amplification (Fig. 6-b), resonance annihilation (Fig. 6-c) and generation of antiresonance/antiphasonance (Fig. 6-d).

#### 3.5.1 Envelope-space diagrams: projections onto the *v-w*_1_ and *v-w*_2_ planes

In analogy to the sinusoidally forced 2D models discussed in Section 3.1.2, we view the dynamics of the oscillatory forced 3D model (30)-(32) as the result of the RLC trajectory tracking the oscillatory motion of the *v*-nullsurface

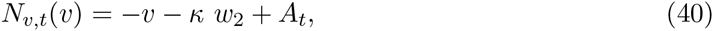

where *A*_*t*_ representes the oscillatory input (29), in between the levels corresponding to *A*_250_ = 1 and *A*_750_ = −1. The *w*_1_- and *w*_2_-nullsurfaces are both given by the same expression

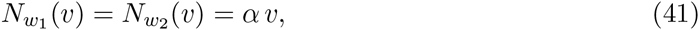

and are independent of the oscillatory input.

In order to facilitate the visualization of this “moving phase-space” we look at its projections onto the *v-w*_1_ and *v-w*_2_ planes, which we refer to simply as the envelope-plane diagrams (Fig. 12). The corresponding *v*-nullclines are given by

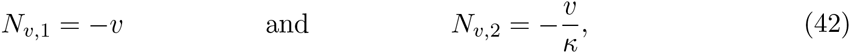

and the associated “moving v-nullclines” are given by

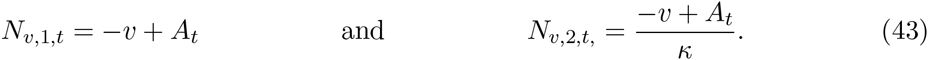

We refer to the projections of the *w*_1_- and *w*_2_-nullsurfaces as the *w*_1_- and *w*_2_-nullclines respetively.

**Figure 12:**
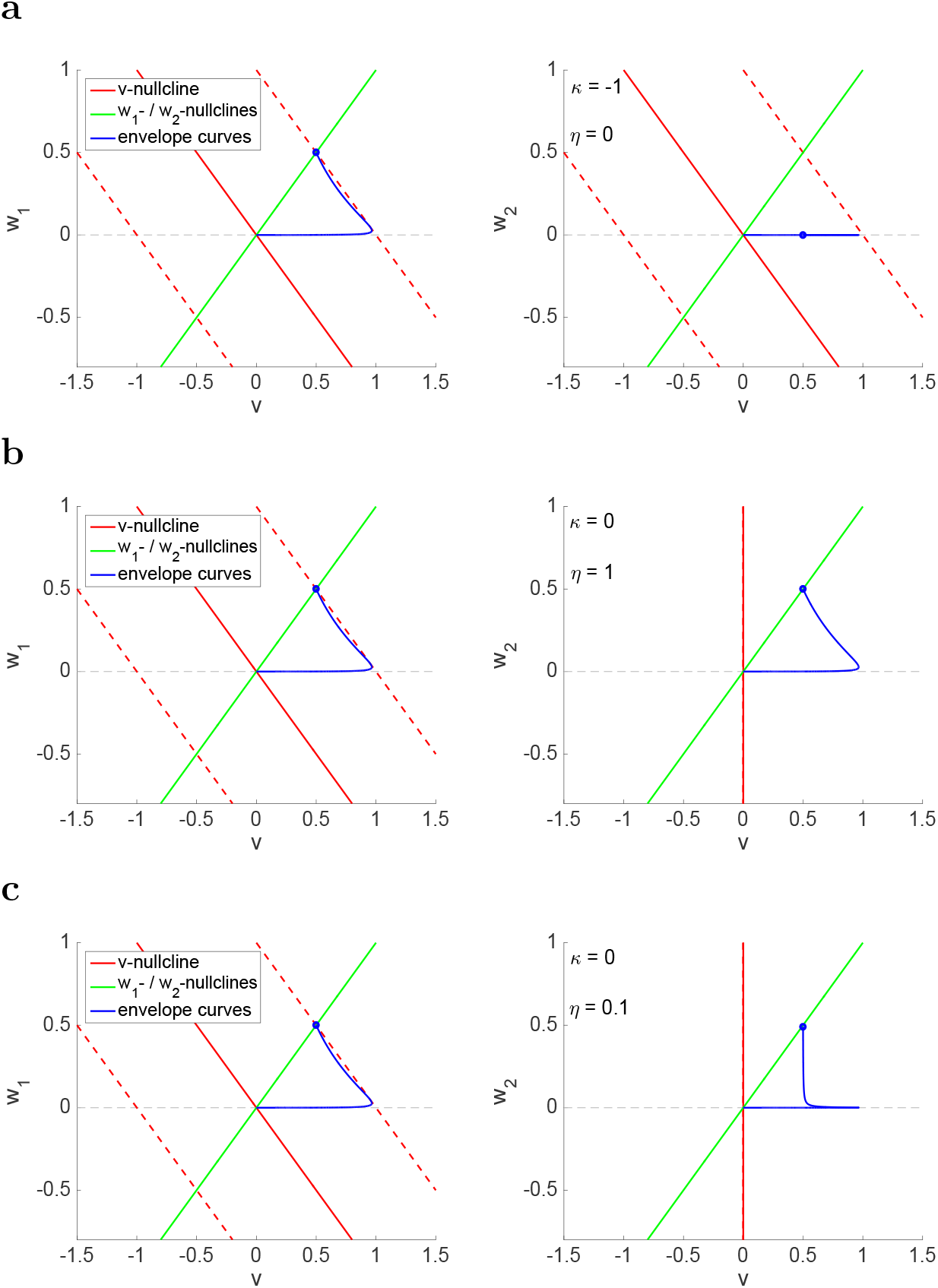
**Envelope-plane diagrams for the linear 3D model in the limiting 2D regimes.** The red and green nullclines are the projections of the *v*-, *w*_1_- and *w*_2_-nullsurfaces onto the *v-w*_1_ (left) and *v-w*_2_ (right) planes. The *v*-nullcline (solid-red) moves cyclically in between the displaced *v*-nullclines for ±*A*_*in*_ (dashed-red) following the sinusoidal input. The (upper) envelope curves (solid blue) join the maximum points of the projection of the limit cycle response trajectories onto the corresponding planes as the input frequency *f* increases from *f* = 0 (intersection between the dashed-red and green lines in the left panel) to *f* → ∞ (intersection between the solid-red and green lines determining the fixed-point in the left panel). The lower envelope curves (not shown) are symmetric to the upper ones with respect to the origin (fixed-point of the autonomous system). **(a)** *κ* = −1 and *η* = 0. *w*_2_ is constant and therefore has no frequency-dependent effect on the 3D system. **(b)** *κ* = 0 and *η* = 1. *w*_2_ is decoupled from the 3D system. **(b)** *κ* = 0 and *η* = 0.1. *w*_2_ is decoupled from the 3D system. We used the following parameter values: *α* = 1 and *ϵ* = 0.05.

We use the limiting 2D regimes presented in Fig. 12 as the baseline for our discussion. The *w*_1_- envelope-plane diagrams shown in Fig. 13 (left) are extensions to 2D type shown in Fig. 3 (panels d and e) for the same parameter values as in Fig. Fig. 12-c. For *f* = 0 (blue dot) the envelope-curve (blue) is located at the intersection of the *v*- and *w*_1_-nullclines, which reflects the fast dynamics of the RLC trajectory along the *w*_1_-nullcline. As *f* →∞ the RLC trajectory approaches the origin. The points in the envelope curve for *f*_*res*_ and *f*_*phas*_ are the peak in the *v*-direction (*f*_*res*_) and the tangency point between the envelope-curve and the dashed-red line (*f*_*phas*_), reflecting the fact that the RLC trajectory and the *v*-nullcline reached their maximum levels at the same time. For *η* = 0 (Fig. 12-a) *w*_2_ is a constant and therefore the envelope-curve (right panel) evolves along the *v*-direction and has no frequency-dependent effect on the system. For *η* = 1 and *κ* = 0 (Fig. 12-b) *w*_2_ and *w*_1_ have identical dynamics and therefore the *w*_1_- and *w*_2_ envelope curves are identical. However, since *κ* = 0, *w*_2_ is following *v*, but not feeding back onto its dynamics. According to eqs. (42) and (43) the *v*-nullcline in the right panel is vertical for all values of *A*_*t*_.

**Figure 13:**
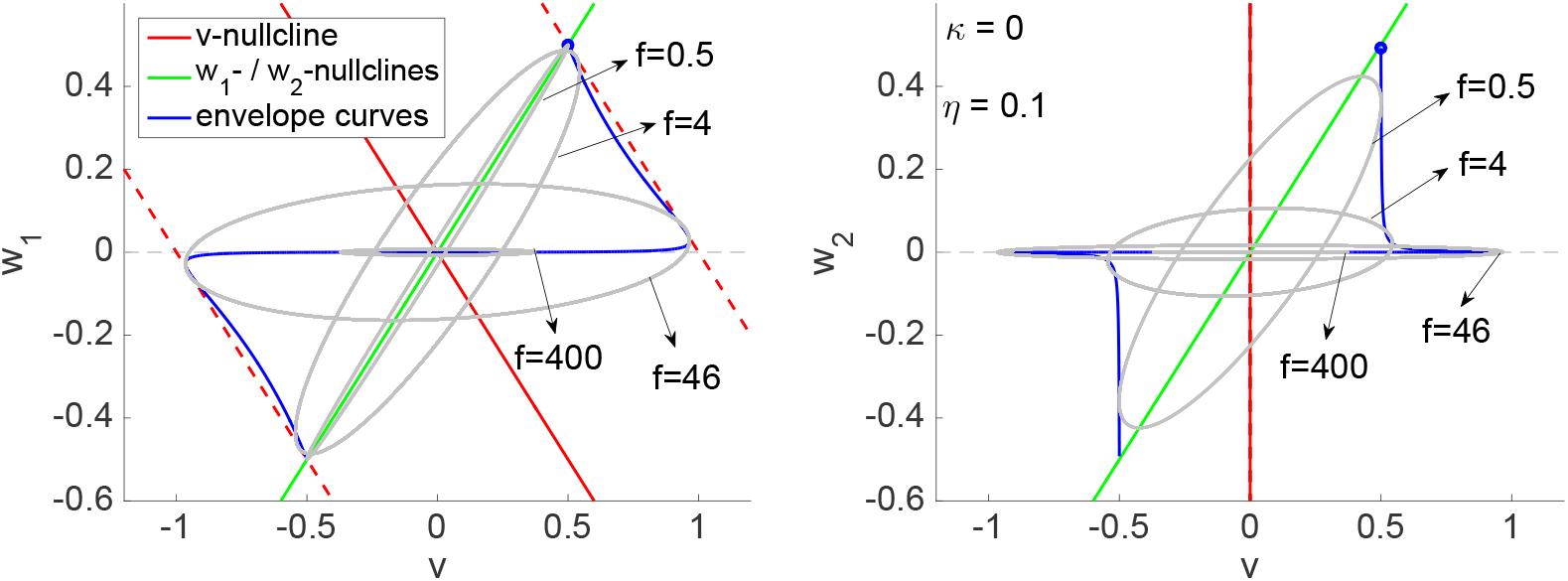
**Projections of the envelope-space diagram onto the** *v-w*_1_ **and** *v-w*_2_ **planes and representative response limit cycle trajectories for the linear 3D model in the limiting 2D regimes.** The paramter values correspond to Fig. 12-c.

In both cases (Figs. 12-a and -b), the evolution of the RLC trajectories with increasing values of *f* is as in Fig. 3-d. For *f* = 0 the RLC trajectory moves along the *w*_1_-nullcline. As *f* increases the RLC trajectory becomes elliptic-like and the major axis rotates towards the *v*-(horizontal) direction as its length increases. As *f* continues to increase, the RLC trajectory begins to shrink and eventually shrinks to the fixed-point (origin) as *f* → ∞. It is important to note that the rays connecting the origin with the points in the envelope-plane curve are not necessarily the major axis of the corresponding elliptic-like RLC trajectories.

The changes in shape (and the direction of the major axis) of the RLC trajectories as *f* increases result from the changes in the effective speed of the RLC trajectories as the balance between *ϵ* and *f* change in eq. (31). The smaller this quotient, the larger the horizontal component of the direction of motion. We refer the reader to [41] for a detailed discussion.

#### 3.5.2 The envelope-plane diagram and RLC trajectories for *w*_1_ and *w*_2_ have different dependences with their time-scale separation *η*

Fig. 12-c illustrates how the shapes of the envelope-plane curves are affected by the relative speed *η* of *w*_1_ and *w*_2_ in the simplest cases in which the 2D system is effectively decoupled from *w*_2_ (*κ* = 0), but the dynamics of *w*_2_ is dependent on *v*. The evolution of the RLC trajectories in the *v-w*_1_ plane (Fig. 13-a, left) is as described above, but the evolution of the RLC trajectories in the *v-w*_2_ plane follows different “rules” (Fig. 13-c, right), which gives rise to the envelope curves with different shapes than in the left panel. First, for low values of f (e.g., *f* = 0.5) the RLC trajectory is wider for *v-w*_2_ than for *v-w*_1_. This is because the horizontal components of the direction of motion are (locally) larger for *w*_2_ than for *w*_1_ due to the smaller time constant of the corresponding equations. This, in turn causes the RLC trajectories’ major axis to rotate faster for *w*_2_ than for *w*_1_ (compare the RLC trajectories for *f* = 4 in Fig. 13). Second, as *f* increases, the RLC trajectories in the *v*- *w*_2_ plane first shrink (from *f* = 0.5 to *f* = 4), then expand (*f* = 46) and then shrinks again (*f* = 400). This is not only in contrast to the behavior of the RLC trajectories in the *v-w*_1_ for this particular value of *η*, but also for other values (e.g., *η* = 1 in Fig. 12-b).

#### 3.5.3 Resonance attenuation and annihilation by increasing levels of the amplifying gating variable with *η* = 1 result from effective 2D dynamics

In Section 3.3.2 we showed that when *τ*_2_ = *τ*_1_ = 100 (Fig. 6-b) decreasing negative values of *g*_2_ cause first an attenuation of the voltage response and then the annihilation of resonance (see also Fig. 9-a and -d1). Here we demonstrate that this phenomenon is generally valid for comparable time constants as captured by *η* = 1 (*τ*_1_ = *τ*_2_).

Figs. 14-a1, -b1 and -c1 show the envelope-plane diagrams for the 3D linear system for three representative values of *κ* < 0 (decreasing from a to c) and *η* = 1. For *κ* = 0 this system exhibits resonance (Fig. 12-b) that results from the 2D linear mechanisms invesigated in [41] and mentioned above. As *κ* decreases the resistance *Z*(0) increases according to (39) moving the blue dot up along the *w*_1_- and *w*_2_-nullclines in the corresponding envelope plane diagrams. Resonance is attenuated because the vertex of the envelope-plane curve remains almost fixed and therefore the angle between the two portions of the envelope-plane curve increases (Fig. 14-a1). As *κ* continues to increase this angle reaches ninety degrees, thus annihilating resonance (Fig. 14-b1 and -c1).

**Figure 14:**
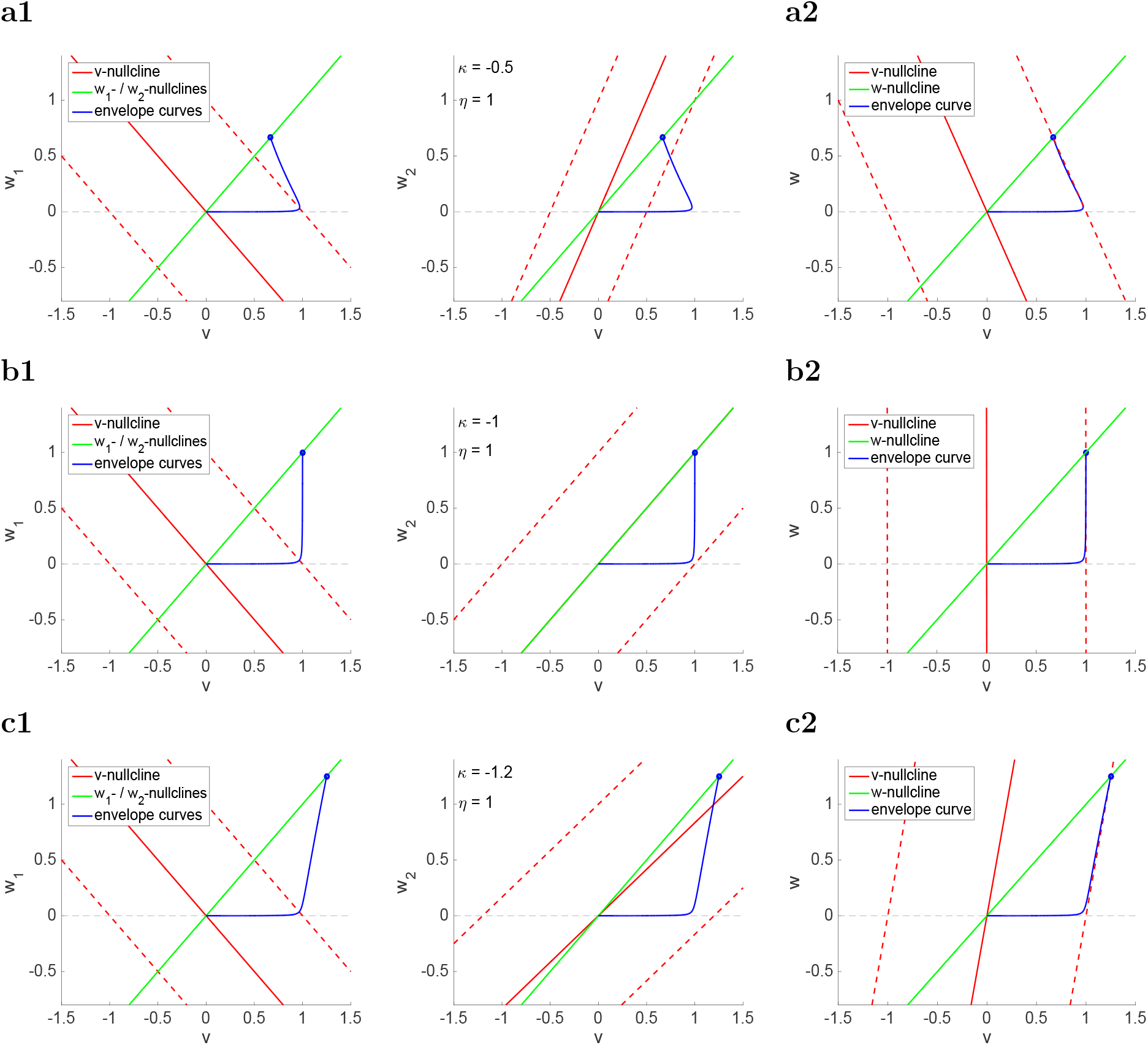
**Projections of the envelope-space diagrams for the linear 3D model with** *η* = 1 (*w*_1_ **and** *w*_2_ **have identical dynamics) on the** *v-w*_1_, *v-w*_2_ **and** *v-w* **planes.** The red and green nullclines are the projections of the *v*-, *w*_1_- and *w*_2_-nullsurfaces onto the *v-w*_1_ (left) and *v-w*_2_ (right) planes, respectively. The *v*-nullcline (solid-red) moves cyclically in between the displaced *v*-nullclines for ±*A*_*in*_ (dashed-red) following the sinusoidal input. The (upper) envelope curves (solid blue) join the maximum points of the projection of the limit cycle response trajectories onto the corresponding planes as the input frequency *f* increases from *f* = 0 (intersection between the dashed-red and green lines in the left panel) to *f* → ∞ (intersection between the solid-red and green lines determining the fixed-point in the left panel). The lower envelope curves (not shown) are symmetric to the upper ones with respect to the origin (fixed-point of the autonomous system). Panels a2, b2 and c2 show the envelope-plane diagram for the equivalent 2D linear system (33)-(34). **(a)** *κ* = −0.5 and *η* = 1. **(b)** *κ* = −1 and *η* = 1. **(b)** *κ* = −1.2 and *η* = 1. We used the following parameter values: *α* = 1 and *ϵ* = 0.05.

Key to this mechanism is the fact that the envelope-curves’ vertices remain almost unchanged with changing values of *κ*. The primary reason for that is Quasi-2D dynamics for *η* = 1 discussed in Section 3.2 (Fig. 4-b) and also illustrated in Fig. 17-b for *κ* = −1.2. More specifically, the two variables *w*_1_ and *w*_2_ are identical, and therefore the system is effectively 2D and governed by eqs. (33)-(34). Figs. 14-a2, -b2 and -c2 show the envelope-plane diagrams for the 2D system and the same values of *κ* as in the left (a1, b1 and c1) panels. The envelope curves in the 2D *v-w* planes and the 3D projections onto the *v-w*_1_ and *v-w*_2_ planes are almost identical.

As explained in [41], for 2D linear systems the envelope-plane curve remains inside the triangle bounded by the *w*-nullcline, the horizontal *v*-axis and the displaced *v*-nullcline (dashed-red). The first two are independent of *κ*, while the latter experiences a change in slope from negative (Fig. 14-a2) to positive (Fig. 14-c2) as *κ* increases, but not the *v*-intercept, thus preventing the envelope-plane curve to peak at higher values of *v*.

#### 3.5.4 Phasonance annihilation

Phasonance is annihilated for the same reason since as *κ* decreases, the envelope-plane curve looses its ability to “touch” the displaced (dashed) *v*-nullcline and therefore the only tangent point between the envelope-curve and the displaced *v*-nullcline is the blue dot.

#### 3.5.5 Generation of antiresonance and antiphasonance

In Sections 3.3.4 and 3.4.4 we have shown that antiresonance and antiphasonance emerge as the result of the interplay of a slow resonant gating variable (*g*_1_ > 0) and a slower amplifying (*g*_2_ < 0) gating variable (see Figs. 6-d and 9-d2). In terms of the geometric rescaling (Section 2.3.2), these differences in the signs of the linearized conductances and the time constants associated to *w*_1_ and *w*_2_ are given by *κ* < 0 and *η* < 1.

Here we explain these phenomena in terms of the envelope-space diagrams (and their projections onto the *v-w*_1_ and *v-w*_2_ planes). Our results are presented in Figs. 15 and 16. In Fig. 15 we present the *w*_1_- and *w*_2_-envelope-plane diagrams for *η* = 0.1 and the same representative values of *κ* < 0 as in Fig. 14.

**Figure 15:**
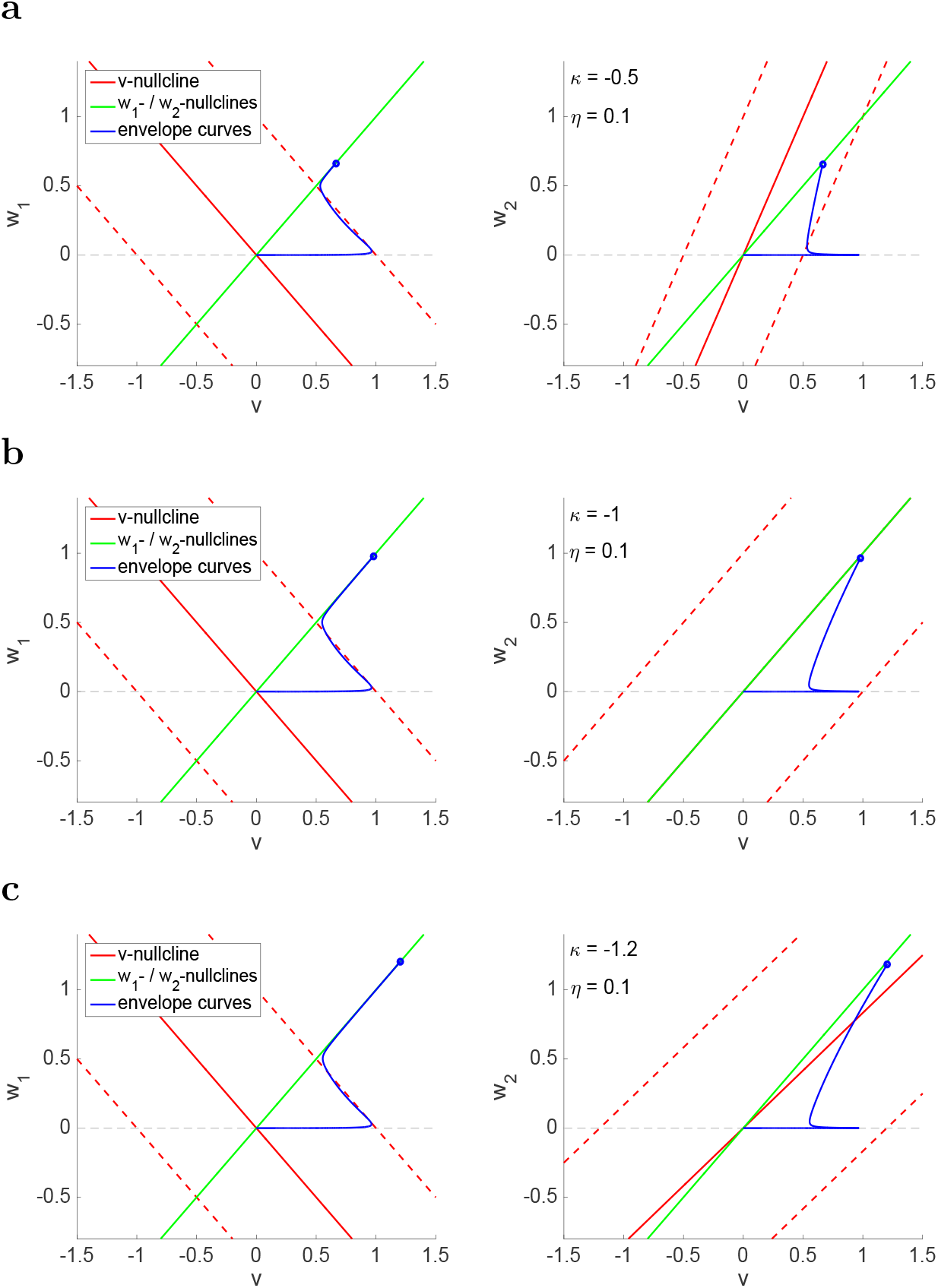
**Projections of the envelope-space diagrams for the linear 3D model onto the** *v-w*_1_ **and** *v-w*_2_ **planes in the limiting 2D regimes.** The red and green nullclines are the projections of the *v*-, *w*_1_- and *w*_2_-nullsurfaces onto the *v-w*_1_ (left) and *v-w*_2_ (right) planes. The *v*-nullcline (solid-red) moves cyclically in between the displaced *v*-nullclines for ±*A*_*in*_ (dashed-red) following the sinusoidal input. The (upper) envelope curves (solid blue) join the maximum points of the projection of the limit cycle response trajectories onto the corresponding planes as the input frequency *f* increases from *f* = 0 (intersection between the dashed-red and green lines in the left panel) to *f* → ∞ (intersection between the solid-red and green lines determining the fixed-point in the left panel). The lower envelope curves (not shown) are symmetric to the upper ones with respect to the origin (fixed-point of the autonomous system). **(a)** *κ* = −0.5 and *η* = 0.1. **(b)** *κ* = −1 and *η* = 0.1. **(b)** *κ* = −1.2 and *η* = 0.1. We used the following parameter values: *α* = 1 and *ϵ* = 0. 05.

As discussed in Section 3.2 (Figs. 4-c and -d for *κ* > 0) for low enough values of *η* < 1 the dynamics of the 3D system is quasi-2D (*v-w*_1_ plane) except for a boundary frequency layer in the vicinity of zero. This remains true for *κ* < 0 (Figs. 17-c and -d). Because of that, not only the horizontal portion of the envelope-curve remains almost unchanged as *κ* changes, but also a significant portion of the envelope-curve for the lower values of *f* along the displaced *v*-nullcline (dashed-red). In contrast, as for the *η* = 1 case explained above, decreasing values of *κ* cause *Z*(0) to move up along the *w*_1_- and *w*_2_-nullclines.

Both antiresonance and antiphasonance are generated because the continuity of the solutions for changing values of *f* requires the envelope curve to decrease from *Z*(0) to the *v-w* − 1 plane at the level of the displaced *v*-nullcline (dashed-red) and then move along this *v*-nullcline towards the vertex of the triangular region. In doing so, *Z*(*f*) reaches a minimum for *f*_*ares*_ > 0 before arriving to the maximum at the vertex (*f* = *f*_*res*_) and *ϕ* “touches” the displaced *v*-nullcline twice: at the crossing point (*f*_*aphas*_) on its way down and at the tangency point (*f*_*phas*_). The behavior of the RLC trajectories for *κ* = −1.2 for representative values of *f* is shown in Fig. 16.

**Figure 16:**
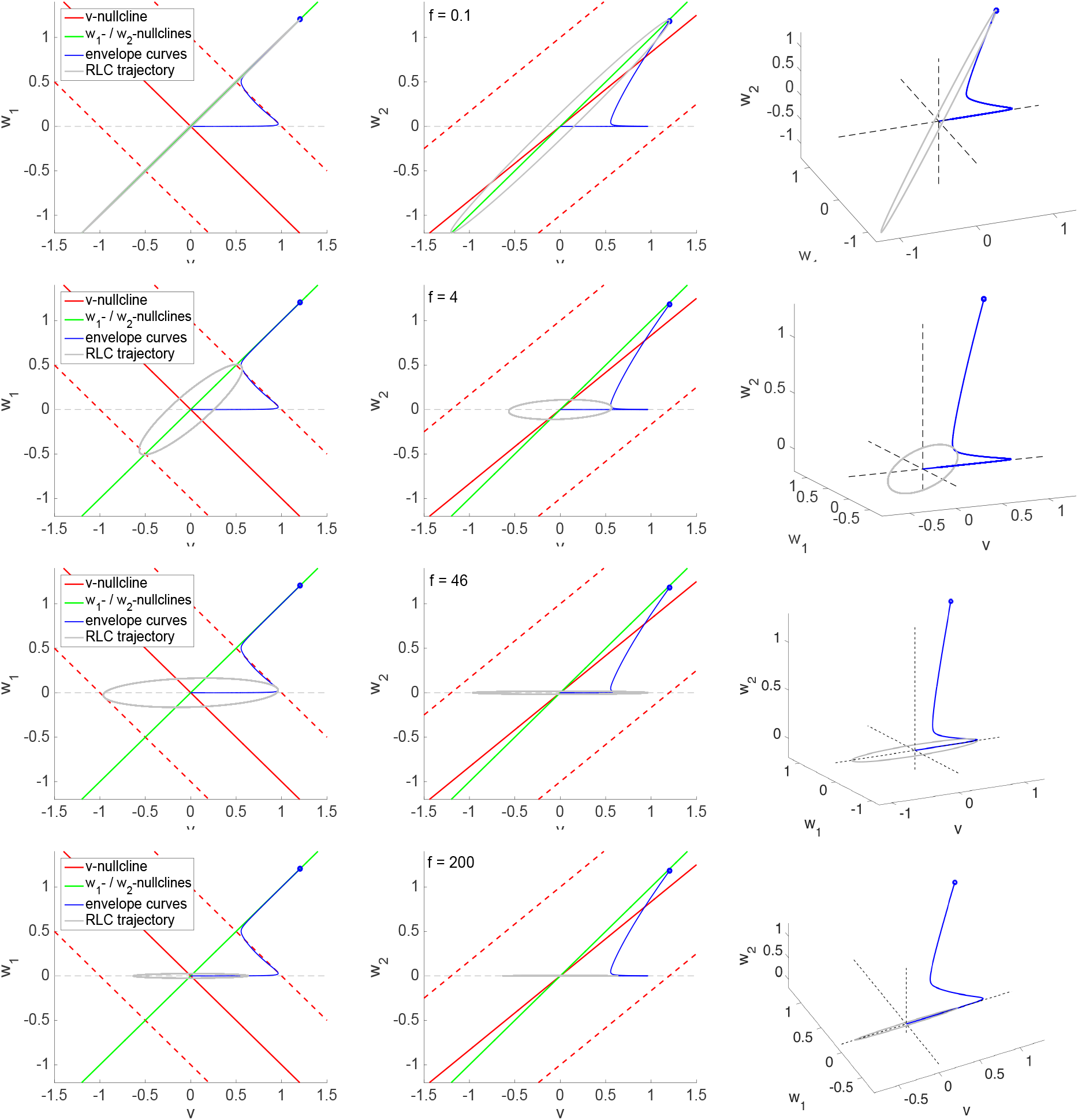
**Projections of the envelope-space diagrams onto the** *v-w*_1_ **and** *v-w*_2_ **planes (left and middle) and representative response limit cycle trajectories (right) for the linear 3D model in an antiresonance regime.** The parameter values correspond to Fig. 15-c. The left and middle panels are the envelope-plane diagrams; i.e., the projections of the 3D envelope-space diagram onto the *v-w*_1_ and *v-w*_2_ planes with the superimposed projections of the RLC trajectories. The right panels show the 3D envelope-curve and the corresponding RLC trajectories.

**Figure 17:**
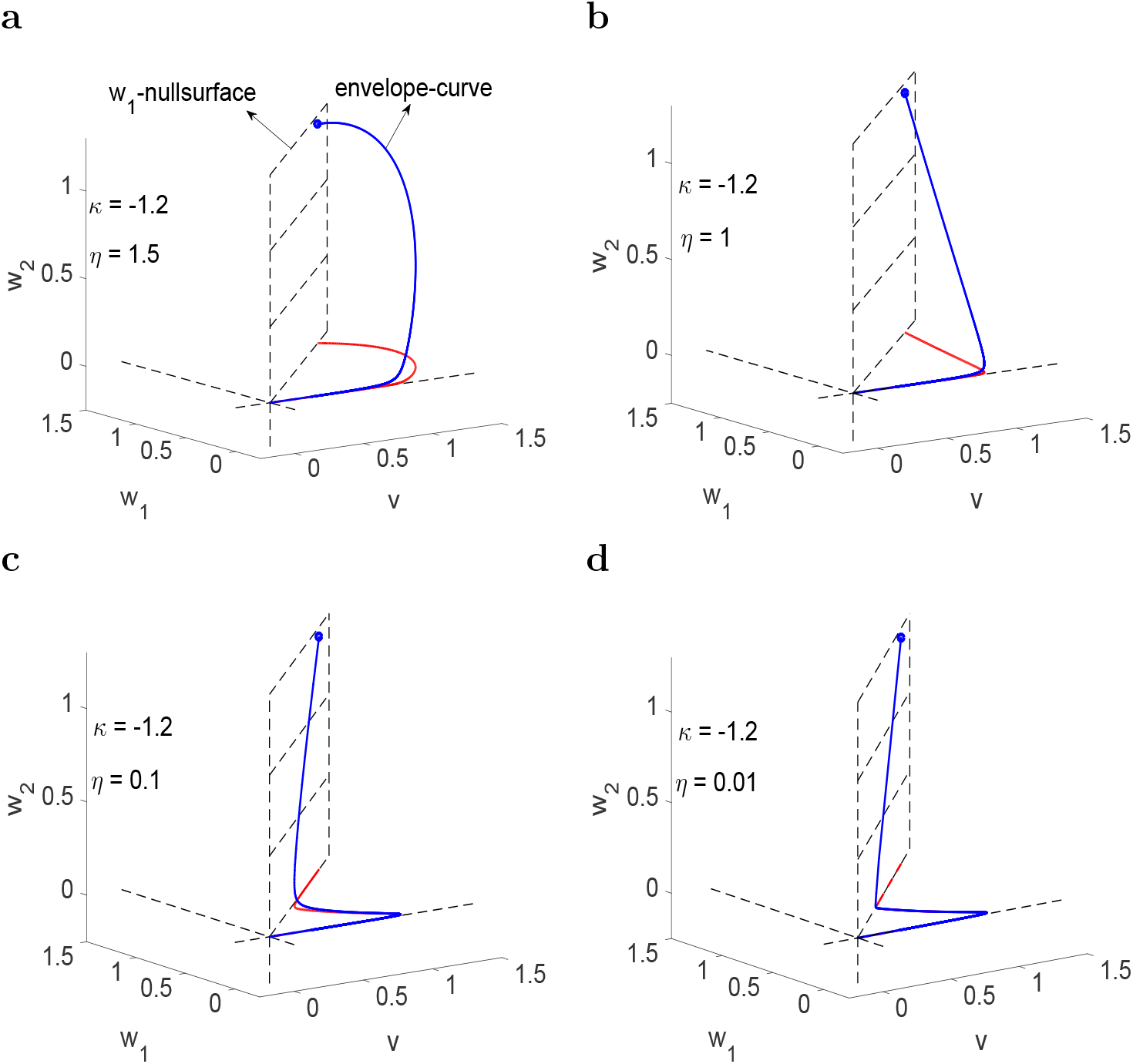
**Envelope-space diagrams for the linear 3D model: resonance modulation, annihilation and generation of antiresonance**. The (upper) envelope curves (solid blue) join the maximum points of the limit cycle response trajectories as the input frequency *f* increases from *f* = 0 (blue dot on the *w*_1_-nullsurface) to *f* → ∞ (at the origin). The red curves are the projection of the envelope-curves onto the *v-w*_1_ (horizontal) plane. The lower envelope curves (not shown) are symmetric to the upper ones with respect to the origin (fixed-point of the autonomous system). We used the following parameter values: *α* = 1, *ϵ* = 0.05 and *κ* = −1.2. The dynamics become increasingly 2D as *η* decreases as indicated by the larger portions of the envelope-curves (blue) that are almost superimposed to their projections (red) onto the *v-w*_1_ planes. **(a)** *η* = 1.5. Resonance is attenuated as *η* decreases. **(b)** *η* = 1. Resonance annihilation. **(c)** *η* = 0.1 Antiresonance and resonance **(d)** *η* = 0.01 Antiresonance and resonance.

## 4 Discussion

In previous work we have investigated the biophysical and dynamic mechanisms of generation of subthreshold resonance and phasonance in 2D linear and linearized conductance-based models [3, 41] and in 2D conductance-based models with nonlinearities of quadratic type [42]. Resonance and phasonance in these models result from the interaction between positive and negative feedback effects provided by the passive (leak) current, a fast amplifying current (slaved to voltage) and a slower resonant current. Each of these currents was assumed to have a single gating variable, although the results are generally valid, at least at the linearized level. More recently, we have investigated how the interplay of positive and negative feedback effects with different time constants shape the intrinsic subthreshold oscillatory patterns in 3D linear and linearized conductance-based models having either two resonant gating variables or one resonant and one amplifying gating variables where the latter is not slaved to voltage and can be slower than the former [50]. These models may also include fast gating variables with instantaneous dynamics (slaved to voltage).

In this paper we have extended this work to include the response of 3D linear models to oscillatory inputs. Our goal was to understand how the cell’s response patterns are shaped by the cooperative activity of the intrinsic feedback processes and time scales and the time scale of the external input determined by its frequency. These models have been studied previously in [2]. One lesson from these and previous studies, including our own, is that the frequency preference properties of cells in response to oscillatory inputs cannot necessarily be predicted from the cell’s intrinsic oscillatory dynamics.

In addition to resonance and phasonance, which are also exhibited by the 2D models, the 3D models exhibit antiresonance and antiphasonance that require the extra degree of freedom provided the third gating variable. To our knowledge this is the first study that addressed the mechanism of generation of antiresonance and antiphasonance. For these phenomena to occur one needs a resonant gating variable (*x*_1_) with slow dynamics, so that the underlying 2D model exhibits resonance, and an amplifying gating variable (*x*_2_) having both slower dynamics (*τ*_2_ > *τ*_1_). The stronger the linearized conductance *g*_2_, the smaller the time constant *τ*_2_ needed for these phenomena to occur (Fig. 11-a and -b). The occurrence of antiphasonance has similar requirements. In fact, there is a complex interplay of the magnitudes and effective time scales of the feedback processes that are responsible for either only modulating the voltage response of the underlying 2D system or creating the more complex types of impedance and phase profiles corresponding to antiresonance and antiphasonance.

Antiresonance has been experimentally reported in hippocampal interneurons [18] and, more recently, in the pyloric network LP neurons (crab stomatogastric ganglion) [37], and it has been shown to be present in the models discussed in [2]. To our knowledge, antiphasonance has not been experimentally reported yet. Our results predict that antiphasonance should occur under similar, but not identical, conditions as antiresonance. Testing the predictions of this study for the presence of both resonance and phasonance in biological neurons involves the manipulation of one or more ionic currents, including maximal conductances and time constants, and can be experimentally performed using the dynamic clamp technique [53–56]. Of particular relevance would be the determination of the differential roles played by different types of amplifying currents such as *I*_*Nap*_ (depolarization-activated) and *I*_*Kir*_ (hyperpolarization-activated) with slow dynamics in biological neurons having different types of resonant currents.

The impedance and phase profiles for linear systems can be computed analytically (see Appendix). These expressions are useful to generate graphs. However, because of their complexity, they provide little information and almost no insight into the dynamic mechanisms leading to the preferred response patterns in both amplitude and phase, including the roles played by the different effective time scales and their complex interaction. To address these issues we ave extended the envelope-space diagram approach introduced in [41, 42] to include the third variable, and we have constructed the envelope curves and their projections onto the appropriate planes. In addition to aid in the mechanistic analysis of the 3D linear system, this approach can be extended to the investigation of the preferred frequency responses to oscillatory inputs of nonlinear 3D systems where the analytic calculations are not possible.

Following previous work [41, 42] we rescaled time by the input frequency *f* in such a way that the sinusoidal input has frequency one for all values of *f*, which instead affect the speed of the voltage response (trajectories in the phase-space), but not the direction of the underlying vector field. By transforming *f* into a time constant factor we were able to capture the different balances that arise between *f* and the cell’s intrinsic time constants as *f* changes. This, in turn, was functional in identifying the mechanism of generation of antiresonance and antiphasonance as an asymptotic boundary layer problem in the frequency domain. For large enough values of *f*, away from *f* = 0, the dynamics are quasi-2D and the impedance profile decreases as *f* decreases towards the boundary layer. In contrast, the resistance is controlled by the 3D system and is higher than this limit in the quasi-2D regime. The trough in the impedance profile is created because the continuity of the solution requires the impedance profile to connect the portions belonging to the two regimes. If in doing so the phase profile crosses the zero value, then antiphasonance is also generated.

For our simulations on the effects of *g*_2_ and *τ*_2_ on the cell’s response patterns we used a baseline set of parameter values for the underlying 2D system (for *v* and *x*_1_) such that it exhibits resonance in a regime where there are not intrinsic subthreshold oscillations. Our envelope-plane diagram analysis demonstrates that the results of our simulations have a more general validity than the restricted considered scenarios. Additionally, as for the 2D models [41,42], this analysis shows that the mechanism of generation of resonance / phasonance and antiresonance / antiphasonance does not qualitatively depend on whether the fixed-point is a (stable) node or a focus. However, a detailed quantification of the dependence of the attributes of the impedance and phase profiles for the 3D linear system on the model parameters requires more research.

In [3, 42], for 2D conductance-based models we have shown through their linearization (or “quadratization”) that changes in the levels of *I*_*h*_ (resonant, hyperpolarization-activated) and *I*_*M*_ (resonant, depolarization-activated) have opposite effects on the dependence of some of the attributes of the impedance and phase profiles (e.g., *f*_*res*_ and *f*_*phas*_) on their maximal conductances when the levels of *I*_*Nap*_ were high enough, while these effects are qualitatively similar for the two resonant currents when the levels of *I*_*Nap*_ were lower. We hypothesize that these features of 2D models will also be present in 3D models and will extend to the antiresonance / antiphasonance attributes. Additional research is needed to test this both theoretically and experimentally and to determine whether additional differential properties emerge between the different types of resonant and amplifying currents.

Antiresonance can be viewed as a neuron’s “anti-preferred” frequency response: a frequency selected by the neuron at which the voltage response is depressed or minimized. Our results indicate an alternative, but complementary view according to which the generation of antiresonance (*Q*_*z*_ > *Q*_0_) can operate as an additional mechanism of amplification of the voltage response in the resonant frequency band where both *Z*_*max*_ and *Z*_0_ may remain unchanged as certain control parameter changes.

Our results open several questions about the neuron’s response properties to oscillatory input. Further research is needed to extend these results to include the effects of the model nonlinearities, and the consequences of the presence of antiresonance and antiphasonance for spiking dynamics and network oscillations.

## Acknowledgments

This work was supported by the NSF grant DMS-1313861 (HGR).

## A Impedance and phase profiles for 2D and 3D linear systems: Analytic expressions

In order to analytically compute he impedance and phase profiles for 3D linear generic systems we use

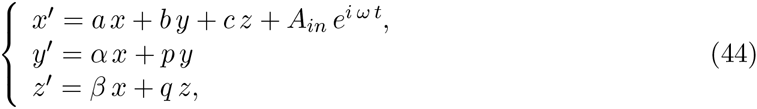

where *a, b, c, α, β, p* and *q* are constant, *ω* > 0 and *A*_*in*_ ≥ 0.

The characteristic polynomial for the corresponding homogeneous system (*A*_*in*_ = 0) is given by

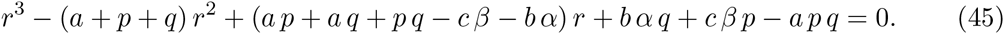

The particular solution to system (44) has the form

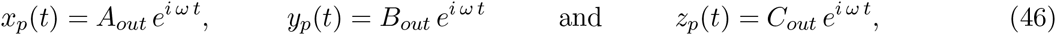

Substituting (46) into system (44), rearranging terms, and solving the resulting algebraic system one obtains

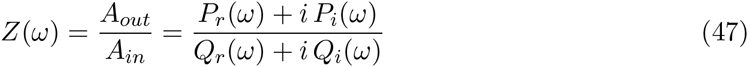

where

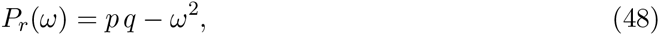

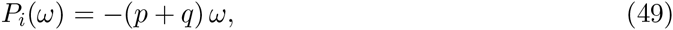

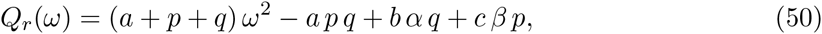

and

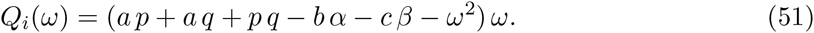

From (47)

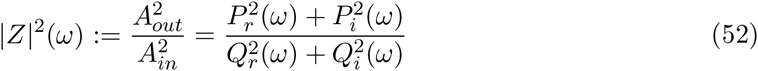

and

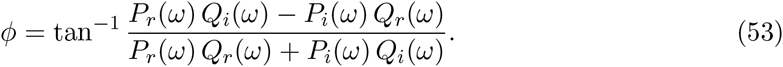

For a 2D linear system (*c* = *q* = 0), the characteristic polynomial for the corresponding homogeneous system (*A*_*in*_ = 0) is given by

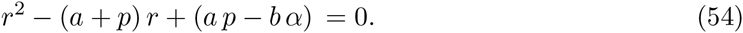

The roots of the characteristic polynomial are given by

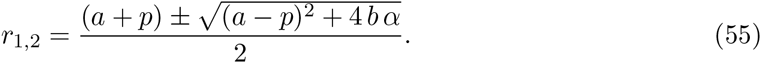

From eq. (55), the homogeneous (unforced) system displays oscillatory solutions with a natural frequency *f*_*nat*_ (Hz) given by

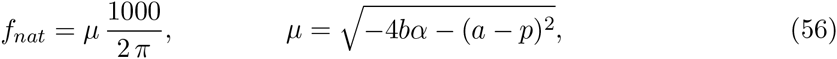

provided 4 *bα* + (*a* − *p*)^2^ < 0.

The impedance amplitude and phase are given, respectively, by

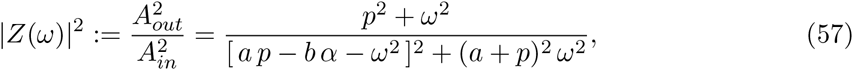

and

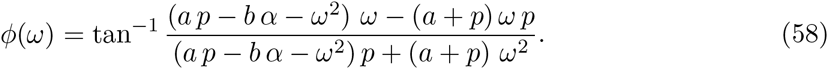

